# Techniques to Produce and Evaluate Realistic Multivariate Synthetic Data

**DOI:** 10.1101/2021.10.26.465952

**Authors:** John Heine, Erin E.E. Fowler, Anders Berglund, Michael J. Schell, Steven Eschrich

## Abstract

**Background:** Data modeling in biomedical-healthcare research requires a sufficient sample size for exploration and reproducibility purposes. A small sample size can inhibit model performance evaluations (i.e., the small sample problem).

**Objective:** A synthetic data generation technique addressing the small sample size problem is evaluated. We show: (1) from the space of arbitrarily distributed samples, a subgroup (class) has a latent multivariate *normal characteristic*; (2) synthetic populations (SPs) of *unlimited* size can be generated from this class with univariate kernel density estimation (uKDE) followed by standard normal random variable generation techniques; and (3) samples drawn from these SPs are statistically like their respective samples.

**Methods:** Three samples (n = 667), selected *pseudo-randomly,* were investigated each with 10 input variables (i.e., X). uKDE (optimized with differential evolution) was used to augment the sample size in X (i.e., the input variables). The enhanced sample size was used to construct maps that produced univariate normally distributed variables in Y (mapped input variables). Principal component analysis in Y produced uncorrelated variables in T, where the univariate probability density functions (pdfs) were approximated as normal with specific variances; a given SP in T was generated with normally distributed independent random variables with these specified variances. Reversing each step produced the respective SPs in Y and X. Synthetic samples of the same size were drawn from these SPs for comparisons with their respective samples. Multiple tests were deployed: to assess univariate and multivariate normality; to compare univariate and multivariate pdfs; and to compare covariance matrices.

**Results:** One sample was approximately multivariate normal in X and all samples were approximately multivariate normal in Y, permitting the generation of *unlimited* sized SPs. Uni/multivariate pdf and covariance comparisons (in X, Y and T) showed similarity between samples and synthetic samples.

**Conclusions:** The work shows that a class of multivariate samples has a latent *normal characteristic*; for such samples, our technique is a simplifying mechanism that offers an approximate solution to the small sample problem by generating similar synthetic data. Further studies are required to understand this latent normal class, as two samples exhibited this characteristic in the study.

## Introduction

Reliable research in predictive modeling requires sufficient data for exploration and reproducibility purposes. This is especially relevant to biomedical-healthcare research, where data can be limited; although this field is broad, a few examples include risk prediction [1], response to therapy [2] and benign malignant classification [3–5]. Unfortunately, data can be limited for variety of reasons: the study of low-incidence diseases or underserved/underrepresented subpopulations [6]; clinic visitation hesitancy [7, 8]; the inability to share data across facilities [9]; cost of molecular tests; and study timeframes. We, the authors, have worked in biomedical-healthcare research for many years and have experienced this persistent problem over decades.

Multivariate modeling is often exploratory and can decrease model stability in various ways. Here we explain frequent approaches that we have experienced. The process starts by analyzing data from the target population for a variety of goals such as: open-ended analyses by studying many different data characteristics searching for correlations and patterns; subgrouping the dataset; testing hypothesis feasibilities with varying endpoints; exploring multiple hypotheses simultaneously; feature selection; selecting the most suitable model; or estimating model parameters with an optimization procedure. In practice, there are virtually *unlimited* ways to search through a given data sample. Data mining of this sort may not always be viewed in the most positive light [10], but on the other hand it is also the nature of discovery, noting there is often a compromise between these positions. Extensive subgroup analyses can effectively deplete the sample. When this applies, we term it the *small sample problem*. In the *final* stage, the fully specified model (i.e., the model with its parameters fixed) is validated with new data to prove its generalizability. Both the exploration and final stages depend critically on having an adequate sample size.

Determining the adequate sample size in the multivariate setting is a difficult task [11] and has relevance to the small sample problem. Adequate multivariate sample size is a function of both the analysis technique and covariance structure. For example, a multivariate two-sample test with normally distributed data and common covariance, Hotelling’s T^2^, is appropriate when comparing mean vectors [12]. In a broad sense, when the variables under consideration tend to be more highly correlated, the adequate sample size decreases and vice versa. Adequate sample size is a function of the number of free model parameters, which does not necessarily correspond with the number of variables [13]. In ordinary linear regression modeling with d noninteracting variables, there are about d parameters that must be determined. In contrast when taken to the limit, partial least squares regression [14] has roughly d^2^ parameters, and deep neural network architectures have even greater number of parameters, requiring large sample sizes for a given design [15] (see related table in [15] for examples). These modeling techniques illustrate that an adequate sample size under one condition may not be optimal for another. It is our premise that the adequate sample size for a given multivariate prediction problem that allows independent validation deserves more attention beyond *larger sample sizes are better,* especially when normality assumptions do not hold. By hypothesis, a technique that can generate realistic synthetic data will provide benefits in modeling endeavors. Such an approach could be used to augment an inadequate sample size for modeling and validation purposes or to study sample size requirements for a given multivariate covariance structure.

Synthetic data applications in health-related research use a variety of techniques. Some methods are used for generating samples from large populations [16–19]. These approaches include hidden Markov models [18], techniques that reconstruct time series data coupled with sampling the empirical probability density function of the relevant variables [16], and methods that estimate probability density functions (pdfs) from the data, not accounting for variable correlation [19]. Other work used moment matching to generate synthetic data but does not consider relative frequencies in the comparison analysis [20]. Discussions on synthetic data generation techniques indicate that the small sample size condition has received little analytical attention [20, 21].

Our synthetic data technique estimates a multivariate pdf for arbitrarily distributed data, including when normal approximations fail to hold [22]. This initial work, based on multivariate kernel density estimation (mKDE) with unconstrained bandwidths, was illustrated with d = 5 data from mammographic case-control data [22]; synthetic populations (SPs) of arbitrarily large size were generated from samples of limited size. However, mKDE has noted efficiency problems for high dimensionality [23, 24]. As the dimensionality increases, the sample size requirement apparently becomes exceeding large, noting this area is under investigation. Although categorizing a problem as high or low dimensionality may be dependent on many factors, reasonable arguments suggest that *high dimensionality* may be defined as 3 < d ≤ 50, where d is the number of variables considered [25]; in that, density estimators should be able to address this range. Here we let d = 10 so that many of the findings can be presented graphically or reasonably tabulated, and modeling problems in healthcare research can be within this range.

In this report, we present modifications to our method to mitigate the mKDE efficiency problem under specific conditions (latent normality) and address synthetic data generation in relatively higher dimensionality (d = 10). This modified approach decomposes an arbitrarily distributed multivariate problem into multiple univariate KDE (uKDE) problems while characterizing the covariance structure independently. We are evaluating whether this approach can transform an arbitrarily distributed multivariate sample into an approximate multivariate normal form, which we define as a sample with a *latent normal characteristic*. In the universe of arbitrarily distributed samples, there is a multivariate normal subgroup. The technique for generating synthetic data for this normal subgroup is relatively straightforward and well-practiced; by hypothesis, our approach seeks to extend these straightforward techniques to the latent normal class by determining when (or if) it exists. Developing the analytics to detect this condition and then leveraging it to generate synthetic data are essential elements of this work.

## 2. Methods

### 2.1 Overview

Our modified synthetic data generation and analytic techniques have sequential components and many related analyses. Therefore, clear definitions, preliminaries, and a brief outline are given before the details are provided. Justifications are also discussed here when warranted.

*Definitions*: *population* is used to define a hypothetical collection of virtually *unlimited* number of either real or synthetic entities from which samples comprised of observations or realizations may exist or can be drawn. The exception for the use of *population* is when explaining differential evolution (DE) optimization [26] used for uKDE bandwidth determination; the *DE-population* is limited and defined specifically. *Sample* defines a collection of n real observations with d attributes (variables) from the space of possible samples, represented mathematically as a n×d matrix (rows=observations, columns=attributes). Column vectors are designated with lower-case bold letters. For example, individual attributes are referred to as **x**, a column vector. Vector components are designated with lower-case subscripted letters. The components of **x** are referenced as x_j_ for j = 1, 2, …, d and assumed continuous. The multivariate pdf for **x** is p(**x**). X refers to the input variable space, that is, **x** exists in the X representation. We assume p(**x**) exists at the population level, but not accessible. In practice, we evaluated normalized histograms throughout this work for all variables considered both univariate and multivariate (i.e., empirical pdfs), also referred to as pdfs for brevity; we use this term to refer to attributes at the population level as well as at the sample level. One-dimensional (1D) marginal pdfs for p(**x**) are expressed as p_j_(x_j_). Matrices are designated with upper case bold lettering. For example, **X** is the n×d matrix with n observations of **x** in its rows (i.e., the i^th^ row contains the d attributes for the i^th^ observation and the j^th^ column of **x** has n realizations of x_j_). Double subscripts are used for both specific realizations and matrix element indices. That is, x_ij_ is the j^th^ component for the i^th^ realization in X (also is the indexing for **X**). Variables in X are mapped to the Y representation. This creates the corresponding entities in Y: (1) the vector **y** with d components; (2) the multivariate pdf, g(**y**), and its marginal pdfs, g(y_j_); and (3) the matrix **Y** defined analogously as **X**. We also work in the T representation (uncorrelated variables) as explained below, where **t**, t_j_, and **T** are defined similarly. Likewise, r(**t**) is the multivariate pdf in T with marginals, r_j_(t_j_). We define the cumulative probability functions (i.e., the indefinite integral approximation for a given univariate pdf) for p_j_(x_j_) and g_j_(y_j_) as P_j_(x_j_) and G_j_(y_j_), respectively. Covariance quantities are calculated with the normal multivariate form: E(w - m_w_)(v - m_v_), where E is the expectation operator, w and v are arbitrary random variables with means m_w_ and m_v_. The corresponding covariance matrices are expressed as **C**_x_, **C**_y_, and **C**_t_, respectively (or **C**_k_ generically). When an entity is given the subscript, s, it then defines the corresponding synthetic entity. Standardized normal defines a zero mean–unit variance normal pdf, used in both the univariate and multivariate scenarios. *Parametric* refers to functions that can be expressed in closed form.

*Preliminaries*: general biomedical healthcare data characteristics are discussed to overview the data characteristics. Measurements such as body mass index (BMI) and age, or measurements taken from image data can have right-skewed pdfs because they are often positive-valued and not inclusive of zero (see figures 1–3 for examples). Such measures can bear varying levels of correlation (see lower parts of tables 1–3). Thus, an arbitrary p(**x**) may not lend itself to parametric modeling in X (i.e., normality and non-correlation approximations do not apply in many instances). To render such data into a more tractable form, a series of steps (see Figure 4) were taken to condition **X**; by premise, these steps will permit characterizing the sample with parametric means and then generating similar multivariate synthetic data without mKDE.

**Figure 1.**
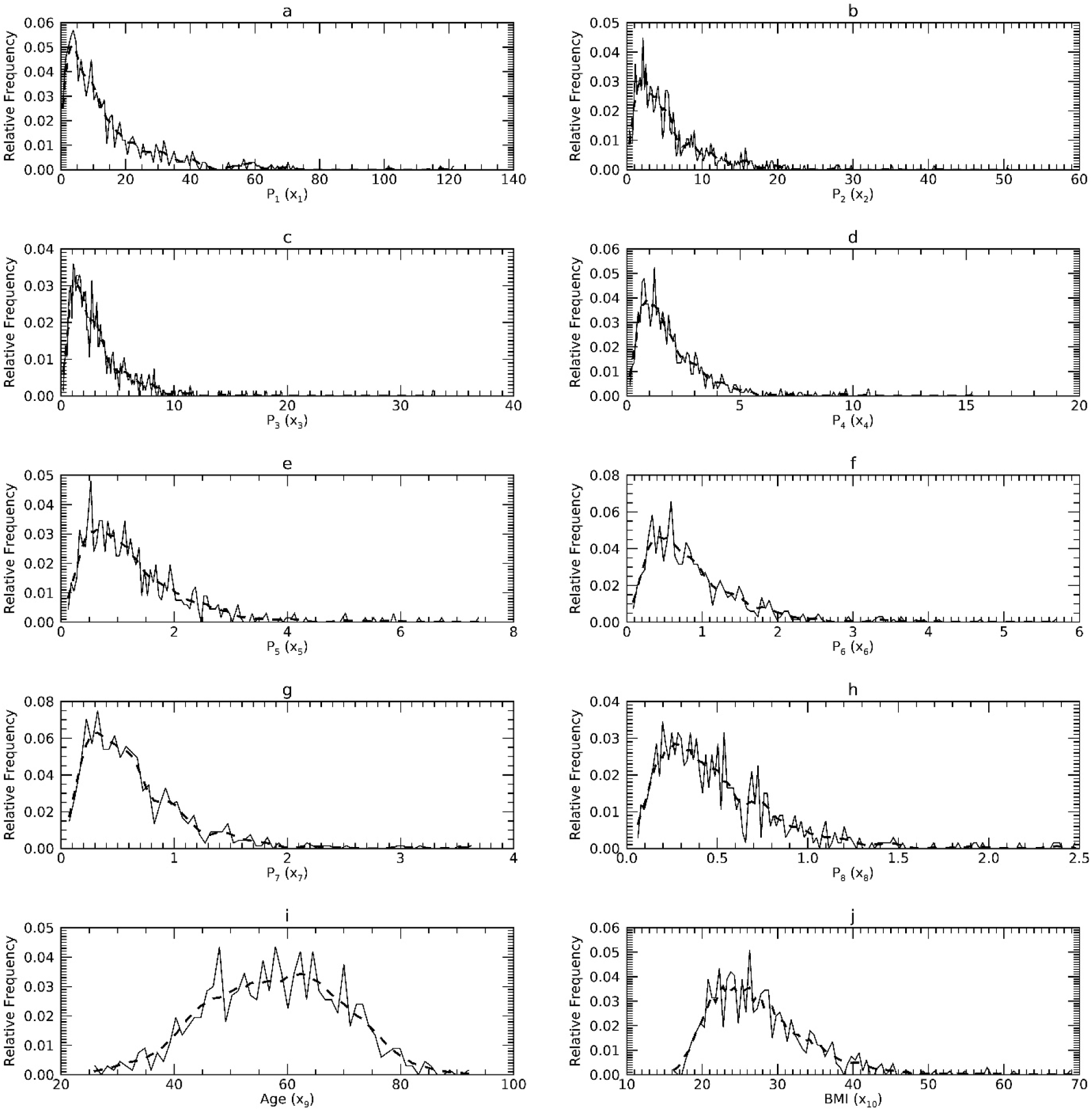
Marginal Probability Density Functions (pdfs) for DS1 in the X representation: each pdf for DS1 (solid) is compared with its corresponding pdf from synthetic data (dashes). The x-axis cites the variable name from its respective resource and its index name parenthetically (x_j_).

**Figure 2.**
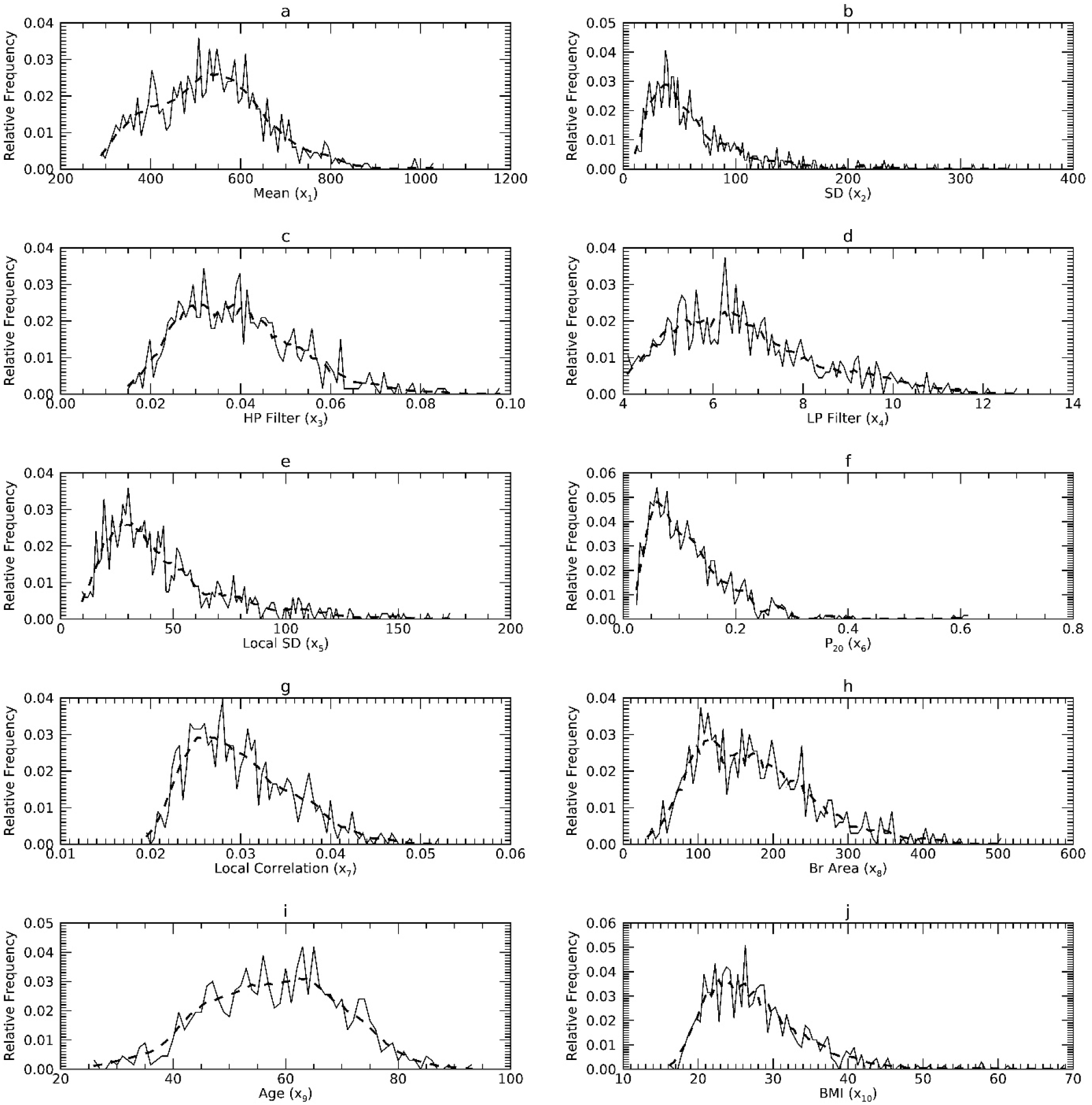
Marginal Probability Density Functions (pdfs) for DS2 in the X representation: each pdf for DS2 (solid) is compared with its corresponding pdf from synthetic data (dashes). The x-axis cites the variable name from its respective resource and its index name parenthetically (x_j_).

**Figure 3.**
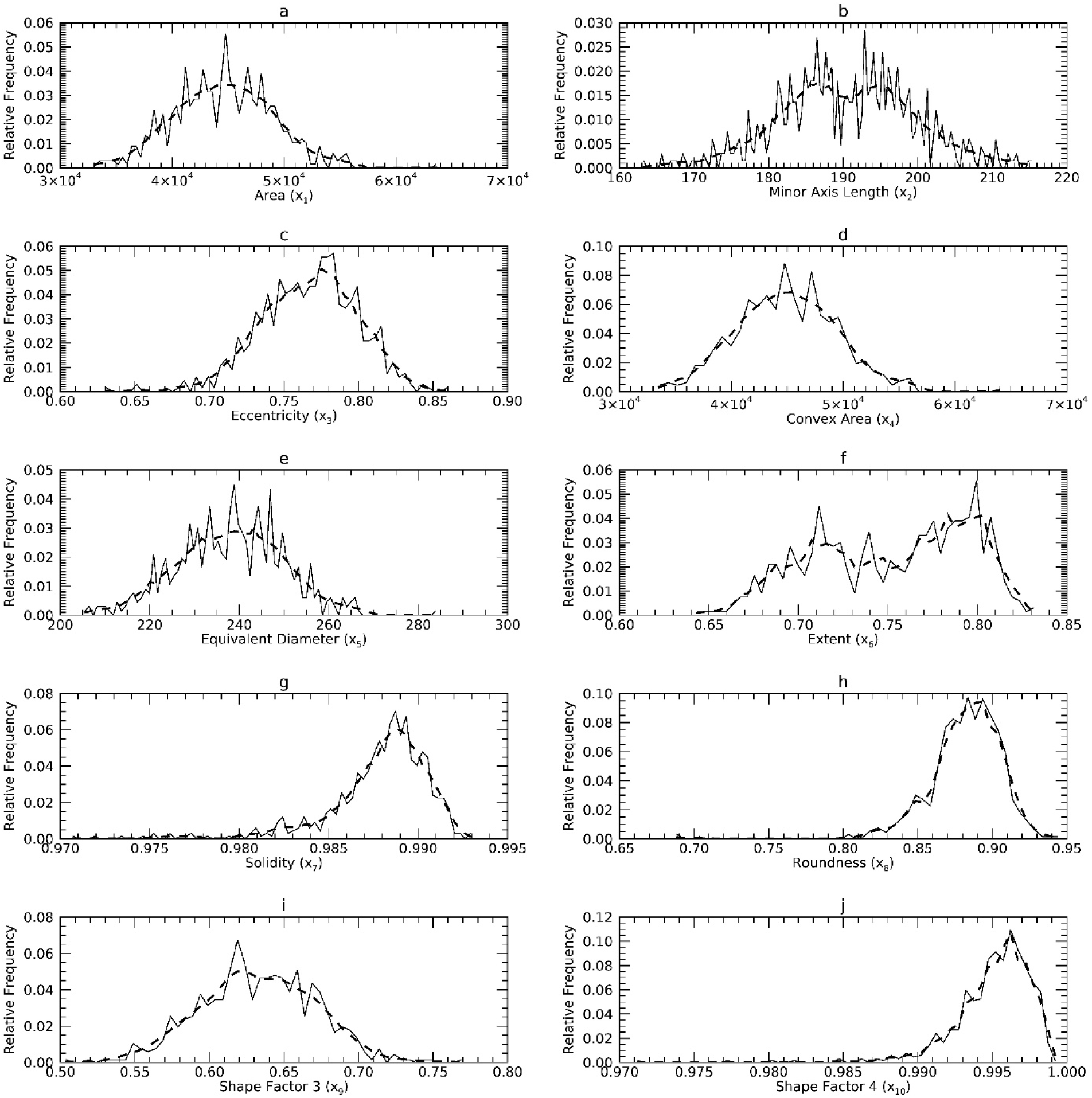
Marginal Probability Density Functions (pdfs) for DS3 in the X representation: each pdf for DS2 (solid) is compared with its corresponding pdf from synthetic data (dashes). The x-axis cites the variable name from its respective dataset and its index name parenthetically (x_j_).

**Figure 4.**
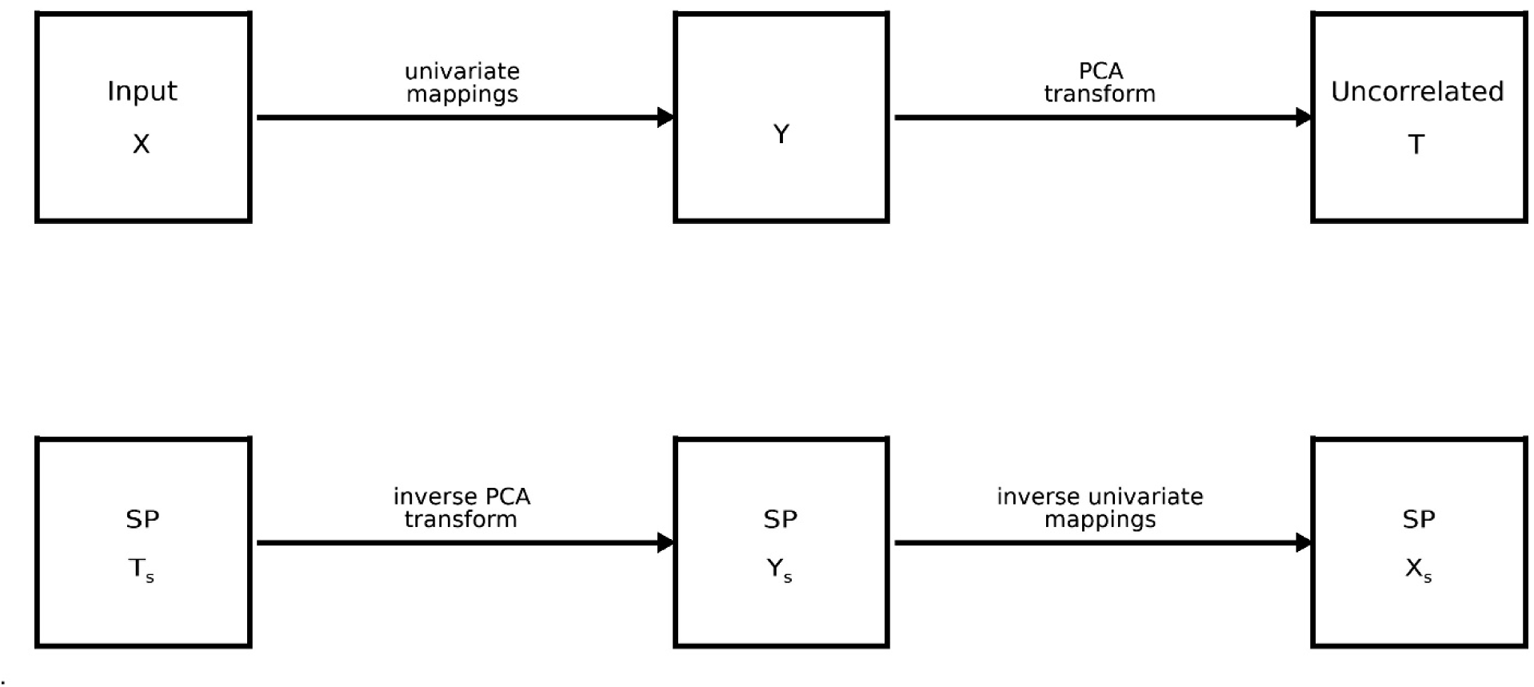
Processing Flow: The top row shows the processing flow for the sample. The reversed processing flow for the SP generation is shown in the bottom row.

**Table 1.** Covariance and Correlation for Dataset 1: In both tables, entries on the diagonal and above give covariance quantities. Entries below the diagonals provide the respective Pearson correlation coefficients. Table 1a gives the X representation quantities and 1b the Y representation quantities. The covariance quantities were generated from the respective sample. Parenthetically, 95% confidence intervals generated from synthetic samples are cited below the respective covariance quantity.

|  | P<sub>1</sub> (x<sub>1</sub>) | P<sub>2</sub> (x<sub>2</sub>) | P<sub>3</sub> (x<sub>3</sub>) | P<sub>4</sub> (x<sub>4</sub>) | P<sub>5</sub> (x<sub>5</sub>) | P<sub>6</sub> (x<sub>6</sub>) | P<sub>7</sub> (x<sub>7</sub>) | P<sub>8</sub> (x<sub>8</sub>) | Age (x<sub>9</sub>) | BMI (x<sub>10</sub>) |
| --- | --- | --- | --- | --- | --- | --- | --- | --- | --- | --- |
| P<sub>1</sub> (x<sub>1</sub>) | 2.27E+02 | 7.27E+01 | 3.61E+01 | 1.97E+01 | 1.18E+01 | 8.16E+00 | 5.76E+00 | 4.27E+00 | -4.13E+01 | -1.98E+01 |
|  | (1.69E+02, 2.88E+02) | (5.38E+01, 9.44E+01) | (2.54E+01, 4.83E+01) | (1.47E+01, 2.58E+01) | (9.17E+00, 1.53E+01) | (6.16E+00, 1.04E+01) | (4.30E+00, 7.19E+00) | (3.21E+00, 5.26E+00) | (-5.15E+01, -2.20E+01) | (-2.59E+01, -1.18E+01) |
| P<sub>2</sub> (x<sub>2</sub>) | 0.9150 | 2.78E+01 | 1.41E+01 | 7.78E+00 | 4.68E+00 | 3.21E+00 | 2.25E+00 | 1.66E+00 | -9.34E+00 | -5.59E+00 |
|  |  | (1.97E+01, 3.62E+01) | (9.42E+00, 1.89E+01) | (5.55E+00, 1.01E+01) | (3.52E+00, 5.96E+00) | (2.35E+00, 4.06E+00) | (1.64E+00, 2.81E+00) | (1.23E+00, 2.05E+00) | (-1.49E+01, -4.84E+00) | (-7.08E+00, -2.12E+00) |
| P<sub>3</sub> (x<sub>3</sub>) | 0.8570 | 0.9600 | 7.82E+00 | 4.30E+00 | 2.58E+00 | 1.77E+00 | 1.22E+00 | 8.95E-01 | -4.22E+00 | -2.29E+00 |
|  |  |  | (4.78E+00, 1.10E+01) | (2.89E+00, 5.73E+00) | (1.87E+00, 3.35E+00) | (1.25E+00, 2.31E+00) | (8.75E-01, 1.59E+00) | (6.61E-01, 1.16E+00) | (-6.83E+00, -1.60E+00) | (-2.92E+00, -2.31E-01) |
| P<sub>4</sub> (x<sub>4</sub>) | 0.8230 | 0.9270 | 0.9660 | 2.53E+00 | 1.52E+00 | 1.04E+00 | 7.18E-01 | 5.32E-01 | -1.65E+00 | -1.33E+00 |
|  |  |  |  | (1.81E+00, 3.28E+00) | (1.15E+00, 1.93E+00) | (7.74E-01, 1.32E+00) | (5.41E-01, 9.14E-01) | (4.09E-01, 6.73E-01) | (-3.25E+00, -3.05E-01) | (-1.63E+00, -9.54E-02) |
| P<sub>5</sub> (x<sub>5</sub>) | 0.8020 | 0.9060 | 0.9410 | 0.9760 | 9.59E-01 | 6.48E-01 | 4.47E-01 | 3.32E-01 | -9.87E-01 | -8.59E-01 |
|  |  |  |  |  | (7.51E-01, 1.20E+00) | (5.03E-01, 8.07E-01) | (3.52E-01, 5.63E-01) | (2.65E-01, 4.17E-01) | (-1.96E+00, -1.37E-01) | (-1.03E+00, -6.19E-02) |
| P<sub>6</sub> (x<sub>6</sub>) | 0.8030 | 0.9040 | 0.9380 | 0.9720 | 0.9810 | 4.54E-01 | 3.12E-01 | 2.33E-01 | -6.05E-01 | -5.83E-01 |
|  |  |  |  |  |  | (3.49E-01, 5.67E-01) | (2.42E-01, 3.90E-01) | (1.84E-01, 2.89E-01) | (-1.23E+00, 2.02E-02) | (-6.93E-01, -2.55E-02) |
| P<sub>7</sub> (x<sub>7</sub>) | 0.8100 | 0.9040 | 0.9230 | 0.9570 | 0.9690 | 0.9820 | 2.22E-01 | 1.65E-01 | -4.09E-01 | -4.28E-01 |
|  |  |  |  |  |  |  | (1.73E-01, 2.78E-01) | (1.30E-01, 2.04E-01) | (-8.82E-01, -8.18E-03) | (-4.93E-01, -2.36E-02) |
| P<sub>8</sub> (x<sub>8</sub>) | 0.8000 | 0.8900 | 0.9050 | 0.9450 | 0.9590 | 0.9750 | 0.9900 | 1.25E-01 | -2.62E-01 | -3.10E-01 |
|  |  |  |  |  |  |  |  | (9.99E-02, 1.54E-01) | (-5.91E-01, 5.92E-02) | (-3.64E-01, -1.00E-02) |
| Age (x<sub>9</sub>) | -0.2330 | -0.1510 | -0.1290 | -0.0884 | -0.0859 | -0.0765 | -0.0739 | -0.0631 | 1.38E+02 | -4.74E+00 |
|  |  |  |  |  |  |  |  |  | (1.30E+02, 1.57E+02) | (-1.09E+01, 7.87E-01) |
| BMI (x<sub>10</sub>) | -0.1950 | -0.1580 | -0.1220 | -0.1240 | -0.1300 | -0.1290 | -0.1350 | -0.1300 | -0.0600 | 4.52E+01 |
|  |  |  |  |  |  |  |  |  |  | (3.72E+01, 5.31E+01) |

**Table 1. Covariance and Correlation for Dataset 1:** In both tables, entries on the diagonal and above give covariance quantities.
|  | P<sub>1</sub> (y<sub>1</sub>) | P<sub>2</sub> (y<sub>2</sub>) | P<sub>3</sub> (y<sub>3</sub>) | P<sub>4</sub> (y<sub>4</sub>) | P<sub>5</sub> (y<sub>5</sub>) | P<sub>6</sub> (y<sub>6</sub>) | P<sub>7</sub> (y<sub>7</sub>) | P<sub>8</sub> (y<sub>8</sub>) | Age (y<sub>9</sub>) | BMI (y<sub>10</sub>) |
| --- | --- | --- | --- | --- | --- | --- | --- | --- | --- | --- |
| P<sub>1</sub> (y<sub>1</sub>) | 1.00E+00 | 9.48E-01 | 9.07E-01 | 8.78E-01 | 8.59E-01 | 8.49E-01 | 8.43E-01 | 8.31E-01 | -2.35E-01 | -2.29E-01 |
|  | (9.03e-01, 1.12e+00) | (8.52e-01, 1.06e+00) | (8.11e-01, 1.02e+00) | (7.85e-01, 9.86e-01) | (7.67e-01, 9.66e-01) | (7.54e-01, 9.56e-01) | (7.47e-01, 9.46e-01) | (7.34e-01, 9.35e-01) | (-3.18e-01, -1.60e-01) | (-3.08e-01, -1.53e-01) |
| P<sub>2</sub> (y<sub>2</sub>) | 0.9480 | 1.00E+00 | 9.71E-01 | 9.49E-01 | 9.34E-01 | 9.22E-01 | 9.16E-01 | 9.03E-01 | -1.78E-01 | -1.54E-01 |
|  |  | (9.01e-01, 1.11e+00) | (8.70e-01, 1.09e+00) | (8.52e-01, 1.06e+00) | (8.38e-01, 1.05e+00) | (8.25e-01, 1.03e+00) | (8.17e-01, 1.03e+00) | (8.08e-01, 1.01e+00) | (-2.56e-01, -1.02e-01) | (-2.33e-01, -8.02e-02) |
| P<sub>3</sub> (y<sub>3</sub>) | 0.9070 | 0.9710 | 1.00E+00 | 9.78E-01 | 9.68E-01 | 9.61E-01 | 9.54E-01 | 9.43E-01 | -1.45E-01 | -1.00E-01 |
|  |  |  | (9.01e-01, 1.12e+00) | (8.79e-01, 1.09e+00) | (8.69e-01, 1.08e+00) | (8.63e-01, 1.07e+00) | (8.55e-01, 1.07e+00) | (8.45e-01, 1.05e+00) | (-2.26e-01, -6.76e-02) | (-1.77e-01, -2.53e-02) |
| P<sub>4</sub> (y<sub>4</sub>) | 0.8780 | 0.9490 | 0.9780 | 1.00E+00 | 9.85E-01 | 9.79E-01 | 9.74E-01 | 9.65E-01 | -1.05E-01 | -9.38E-02 |
|  |  |  |  | (9.01e-01, 1.11e+00) | (8.86e-01, 1.10e+00) | (8.82e-01, 1.09e+00) | (8.75e-01, 1.09e+00) | (8.67e-01, 1.08e+00) | (-1.85e-01, -2.75e-02) | (-1.73e-01, -1.72e-02) |
| P<sub>5</sub> (y<sub>5</sub>) | 0.8590 | 0.9340 | 0.9680 | 0.9850 | 1.00E+00 | 9.86E-01 | 9.82E-01 | 9.74E-01 | -9.71E-02 | -9.27E-02 |
|  |  |  |  |  | (9.00e-01, 1.11e+00) | (8.87e-01, 1.10e+00) | (8.81e-01, 1.09e+00) | (8.73e-01, 1.08e+00) | (-1.76e-01, -2.39e-02) | (-1.73e-01, -1.88e-02) |
| P<sub>6</sub> (y<sub>6</sub>) | 0.8490 | 0.9220 | 0.9610 | 0.9790 | 0.9860 | 1.00E+00 | 9.89E-01 | 9.82E-01 | -8.20E-02 | -8.90E-02 |
|  |  |  |  |  |  | (9.01e-01, 1.11e+00) | (8.89e-01, 1.10e+00) | (8.80e-01, 1.10e+00) | (-1.62e-01, -7.43e-03) | (-1.72e-01, -1.48e-02) |
| P<sub>7</sub> (y<sub>7</sub>) | 0.8430 | 0.9160 | 0.9540 | 0.9740 | 0.9820 | 0.9890 | 1.00E+00 | 9.89E-01 | -8.48E-02 | -9.07E-02 |
|  |  |  |  |  |  |  | (8.95e-01, 1.11e+00) | (8.83e-01, 1.10e+00) | (-1.65e-01, -1.01e-02) | (-1.72e-01, -1.52e-02) |
| P<sub>8</sub> (y<sub>8</sub>) | 0.8310 | 0.9030 | 0.9430 | 0.9650 | 0.9740 | 0.9820 | 0.9890 | 1.00E+00 | -6.30E-02 | -8.84E-02 |
|  |  |  |  |  |  |  |  | (8.96e-01, 1.11e+00) | (-1.43e-01, 8.14e-03) | (-1.68e-01, -9.24e-03) |
| Age (y<sub>9</sub>) | -0.2350 | -0.1780 | -0.1450 | -0.1050 | -0.0971 | -0.0820 | -0.0848 | -0.0630 | 1.00E+00 | -6.64E-02 |
|  |  |  |  |  |  |  |  |  | (8.88e-01, 1.11e+00) | (-1.44e-01, 1.88e-02) |
| BMI (y<sub>10</sub>) | -0.2290 | -0.1540 | -0.1000 | -0.0938 | -0.0927 | -0.0890 | -0.0907 | -0.0884 | -0.0664 | 1.00E+00 |
|  |  |  |  |  |  |  |  |  |  | (8.87e-01, 1.11e+00) |

**Table 2.**
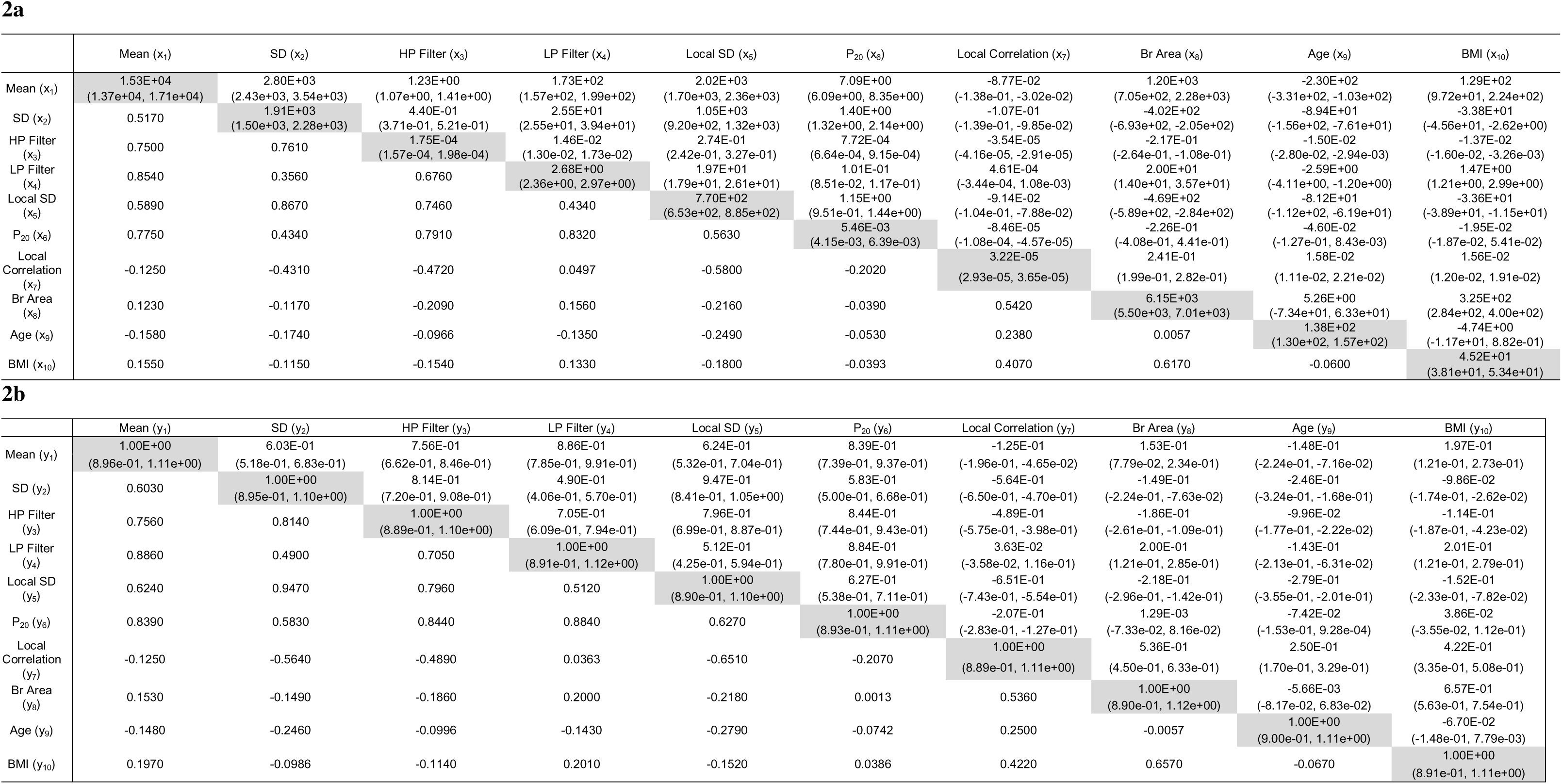
Covariance and Correlation for Dataset 2: In both upper and lower tables, entries on the diagonal and above give covariance quantities. Entries below the diagonals provide the respective Pearson correlation coefficients. Table 2a gives the X representation quantities and 2b the Y representation quantities. The covariance quantities were generated from the respective sample. Parenthetically, 95% confidence intervals generated from synthetic samples are cited below the respective covariance quantity.

**Table 3.**
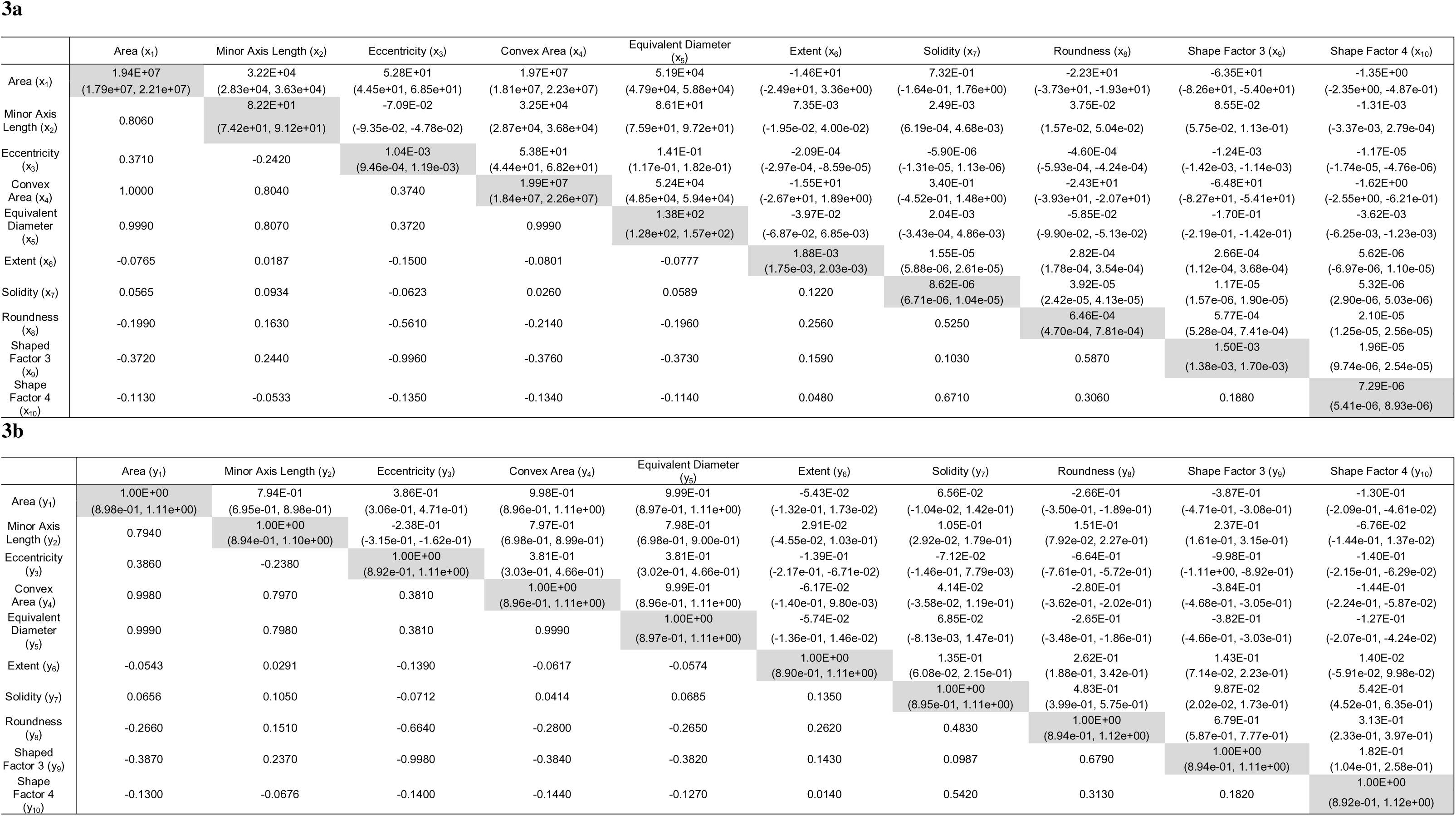
Covariance and Correlation for Dataset 3: In both tables, entries on the diagonal and above give covariance quantities. Entries below the diagonals provide the respective Pearson correlation coefficients. Table 3a gives the X representation quantities and 3b the Y representation quantities. The covariance quantities were generated from the respective sample. Parenthetically, 95% confidence intervals generated from synthetic samples are cited below the respective covariance quantity.

*Outline of the processing steps*: when describing these steps, we also briefly describe the analysis at a given step (also provided in detail in the methods). The process starts with a given sample in X (Figure 4, top left). The processing flow for the sample (X-Y-T) is illustrated in the top row of Figure 4, and the reversed SP generation flow (T_s_-Y_s_-X_s_) in the bottom row.

**Step 1**: Univariate maps were constructed (Figure 4, top-left) to transform a given X measurement to a standardized normal, producing the respective marginal pdf set in Y (Figure 4, top-middle). Maps were constructed with an augmented sample size using optimized uKDE, addressing the small sample size problem.

uKDE was used to generate synthetic x_j_. Here, we augmented the sample size with the goal of filling gaps in the input marginal pdfs of x_j_ (sample) to complement the map constructions. By hypothesis, this step addresses the small sample size problem by guaranteeing continuous smooth maps that will produce standardized normal pdfs from the sample. Synthetic x_j_ generated in this fashion do not maintain the covariance relationships in X and were not used further; only x_j_ from the sample were mapped to Y; KDE was not used beyond this point. There is no guarantee that a set of normal marginals in Y will produce a multivariate normal pdf, although the reverse is always true because a multivariate normal has univariate normal marginals. In practice, g(**y**) from the sample could be assessed at this point to estimate how well it approximates normality. If the latent normal approximation is poor, another synthetic approach could be pursued, or the process could be discontinued. Here we forgo such testing at this step (normality was tested for in steps 3 and 4 instead) and move through all steps to illustrate the techniques. We will also discuss a possible modification that could be investigated when the sample has a poor latent normal characteristic approximation, later in the discussion.

**Step 2**: Principal component analysis (PCA) was used to decouple the variables in Y producing uncorrelated variables in T (Figure 4, top-right).

**Step 3**: Synthetic data was generated in T (Figure 4, bottom-left) as uncorrelated random variables. Here we assumed that each marginal in T from the sample could be approximated as normal with variance given by the j^th^ eigenvalue of **C**_y_. To generate synthetic data, the columns of **T**_s_ were populated as normally distributed random variables with these specified variances (noting, the columns lack correlation). We refer to the realizations in T as the SP (i.e., **T**_s_), noting the column length (number of synthetic entities) can be arbitrarily large. To address the normal characteristic, univariate marginal and multivariate pdfs from the sample were tested for normality at this point.

**Step 4**: The inverse PCA transform (Figure 4, bottom-middle) of **T**_s_ produced the SP in Y (Y_s_), thereby restoring the covariance relationships that were removed in Step 2. We note, this technique (steps 3 and 4) of producing multivariate normal data is a practiced approach when the sample is multivariate normal or well approximated as such. For reference, the multivariate standardized normal in Y is expressed as

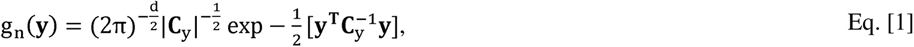

where **C**_y_ can be approximated as the covariance matrix from a given sample. If synthetic samples are poor replicas of their sample, it follows the sample’s g(y) will be a poor approximation of multivariate normality (i.e., the sample is not in the latent normal class).

We evaluated how well the inverse PCA transformation preserved the covariance (C_y_) and the method’s capability of restoring the univariate/multivariate pdfs in Y (i.e., normality comparisons) rather than in Step 2.

**Step 5**: Each synthetic variable in Y is inverse mapped to X (Figure 4, bottom left), which reversed Step 1, thereby producing the SP in X (**X**_s_), and restoring the covariance relationships by hypothesis. The respective pdfs and covariance matrices in X were compared with those from synthetic samples; pdfs were also tested for normality.

It is important to clarify a few aspects of this work. The univariate mapping from X to Y creates a set of univariate marginals normally distributed that can produce a multivariate normal, but not guaranteed. The comparison of univariate marginal pdfs, however, is no guarantee that the respective multivariate pdfs are reasonable facsimiles because the covariance structure has been removed. Many univariate pdf comparisons are provided between samples and synthetic samples in addition to multivariate comparisons, because these allow visualizing similarities with the above stipulations. When a given sample has the latent normal characteristic, the SP generation is greatly simplified, and then it is *defined* by Eq. [1]. The work below shows how to generate synthetic data when this characteristic holds. We use three datasets that were selected *pseudo-randomly*. In the space of samples (virtually unlimited), we do not know the probability that a sample selected at random will have this latent normal characteristic. The main objectives are to present the analysis components with the algorithm for testing for the latent characteristic, give a thorough investigation, and demonstrate that the data synthesis produces realistic data when this latent condition exists, and then discuss further analyses.

### 2.2 Study Data

Samples were derived from two sources of measurements: (1) mammograms and related clinical data (n = 667), and (2) dried beans (n = 13611) [27]. Most technical aspects of these data are not relevant for this report. We used the dried bean data to add variation to the analyses as the variable nomenclatures are very different from the mammogram data, noting at this point the source of data is not germane. From mammograms, we considered two sets of measurements each with d = 10 variables referred to as Sample 1 (DS1) and Sample 2 (DS2). DS1 has 8 double precision measurements from the Fourier power spectrum in addition to age and BMI, both captured as integer variables. The Fourier attributes are from a set of measurements described previously [28]; the first 8 measurements from this set are labeled as P_i_ for i = 1 to 8. The Fourier measures are consecutive and follow an approximate functional form [29], and thus represent components that are very different than those in DS2 (or DS3 below). To cite the covariance quantities and correlation coefficients, we used a modified covariance (covariance for short) table format for efficiency because **C**_k_ is symmetric. In these tables, entries below the diagonal are the respective correlation coefficients, whereas the elements along the diagonal (variance quantities) and above are the covariance quantities. The covariance table for DS1 is shown in Table 1a. DS2 contains 8 double precision summary measurements derived from the image domain: mean, standard deviation (SD); SD of a high-pass (HP) filter output, SD of a low-pass (LP) filter output; local SD summarized; P_20_ Fourier measure (from the set described for DS1 measurements); local spatial correlation summarized [30]; and breast area measured in cm^2^. Age and BMI (from DS1) were also included in this dataset. These variables were selected virtually at random to give d = 10 and possibly provide a different covariance structure than DS1. The covariance table is shown in Table 2a. Neither DS1 nor DS2 were used in our related-prior synthetic data work. Selected measures and realizations from the dried bean dataset [27] are referred to as Sample 3 (DS3). The bean data has 17 measurements (floating point) from 7 bean types. We selected 10 measures at random to make the dimensionality of DS3 compatible with the other two datasets giving this set of variables: area (1); minor axis (5); eccentricity (6); convex area (7); equivalent diameter (8); extent (9); solidarity (10); roundness (11); shape factor 3 (15), and shape factor 4 (16). Here, parenthetical references give the variable number listed in the respective resource (see [27]). Both bean type (bean type = Sira, with n = 2636) and 667 observations were selected at random to create DS3. The covariance table is shown in Table 3a. The analysis of three samples supports the evaluation of the processing scheme under generalized scenarios. The means and standard deviations for x_j_ in each dataset are provided in Table 4. Note, the dynamic range of the means and standard deviations within a given sample vary widely in some instances.

**Table 4.** Mean and standard deviations: This gives the univariate distribution means and standard deviations for all variables by dataset in the X representation.

| DS1 |  |  | DS2 |  |  | DS3 |  |  |
| --- | --- | --- | --- | --- | --- | --- | --- | --- |
| Variable Name | mean | standard deviation | Variable Name | mean | standard deviation | Variable Name | mean | standard deviation |
| P<sub>1</sub> (x<sub>1</sub>) | 1.48E+01 | 1.51E+01 | Mean (x<sub>1</sub>) | 5.35E+02 | 1.24E+02 | Area (x<sub>1</sub>) | 4.48E+04 | 4.41E+03 |
| P<sub>2</sub> (x<sub>2</sub>) | 5.83E+00 | 5.27E+00 | SD (x<sub>2</sub>) | 6.33E+01 | 4.37E+01 | Minor Axis Length (x<sub>2</sub>) | 1.91E+02 | 9.06E+00 |
| P<sub>3</sub> (x<sub>3</sub>) | 3.23E+00 | 2.80E+00 | HP Filter (x<sub>3</sub>) | 4.01E-02 | 1.32E-02 | Eccentricity (x<sub>3</sub>) | 7.69E-01 | 3.23E-02 |
| P<sub>4</sub> (x<sub>4</sub>) | 2.00E+00 | 1.59E+00 | LP Filter (x<sub>4</sub>) | 6.74E+00 | 1.64E+00 | Convex Area (x<sub>4</sub>) | 4.53E+04 | 4.46E+03 |
| P<sub>5</sub> (x<sub>5</sub>) | 1.32E+00 | 9.79E-01 | Local SD (x<sub>5</sub>) | 4.67E+01 | 2.78E+01 | Equivalent Diameter (x<sub>5</sub>) | 2.39E+02 | 1.18E+01 |
| P<sub>6</sub> (x<sub>6</sub>) | 9.21E-01 | 6.74E-01 | P<sub>20</sub> (x<sub>6</sub>) | 1.20E-01 | 7.39E-02 | Extent (x<sub>6</sub>) | 7.58E-01 | 4.33E-02 |
| P<sub>7</sub> (x<sub>7</sub>) | 6.72E-01 | 4.71E-01 | Local Correlation (x<sub>7</sub>) | 3.02E-02 | 5.67E-03 | Solidity (x<sub>7</sub>) | 9.88E-01 | 2.94E-03 |
| P<sub>8</sub> (x<sub>8</sub>) | 5.16E-01 | 3.54E-01 | Br Area (x<sub>8</sub>) | 1.77E+02 | 7.84E+01 | Roundness (x<sub>8</sub>) | 8.84E-01 | 2.54E-02 |
| Age (x<sub>9</sub>) | 5.83E+01 | 1.17E+01 | Age (x<sub>9</sub>) | 5.83E+01 | 1.17E+01 | Shaped Factor 3 (x<sub>9</sub>) | 6.35E-01 | 3.87E-02 |
| BMI (x<sub>10</sub>) | 2.79E+01 | 6.73E+00 | BMI (x<sub>10</sub>) | 2.79E+01 | 6.73E+00 | Shaped Factor 4 (x<sub>10</sub>) | 9.95E-01 | 2.70E-03 |

### 2.3 KDE, optimization, and mapping

The mapping (Step 1) relies on generating synthetic x_j_ with uKDE. Each bandwidth parameter was determined with an optimization process wherein synthetic data from uKDE was compared with the sample. There is a continued feedback loop between the sample, synthetic data generation, and comparison during the optimization process. When the optimization was completed, a given map was constructed.

*uKDE*: for the map construction in Step 1 and as a modification to our previous work [22], uKDE was used to generate realizations from each p_j_(x_j_) given by

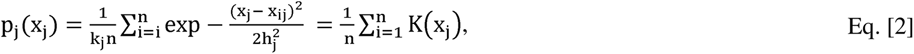

where x_j_ is the synthetic variable for this discussion, x_ij_ are observations from a given sample, k_j_ is a normalization factor, and h_j_ is the univariate bandwidth parameter.

*Optimization*: differential evolution optimization [26] was used to determine each h_j_ in Eq. [2]. The population of candidate h_j_ evolves over generations to a *solution*, as described in detail previously [22]. To form generation zero for a given x_j_, the DE-population (n = 1000) was initialized randomly (uniformly distributed) within these bounds: (0.0, 4 × the variance of x_j_) because a given range should span the solution for h_j_. The DE-population size stays constant across all generations. By expectation, the populations become more fit, according to the fitness function, over generations. A given generation is found by comparing two-DE populations (1000 pair-wise competitions) of candidate h_j_ solutions derived from the previous generation. For a given pairwise competition, two respective SPs were generated for each candidate h_j_. Synthetic samples (n = 667) were drawn from each SP and P_j_(x_j_) from the sample was compared with each P_j_(x_j_) derived from its synthetic sample using Eq. [2]. The h_j_ candidate [used in Eq. (2)] that produced a smaller D_j_ was used to populate the current generation, where D_j_ is the difference metric derived from the fitness function. This process was repeated to obtain 30 generations. In summary, 30×2×1000 synthetic samples were compared with the sample via P_j_(x_j_) comparisons for a given x_j_ to derive its respective h_j_ used in Eq. (2). A given X to Y map was then constructed with x_j_ sampled from Eq. (2) with h_j_ = E[terminal population of candidate h_j_].

As a modification, the Kolmogorov Smirnov (KS) test [31] was used as the fitness function in the DE optimization. This is a nonparametric test that can be used to compare two numerically derived cumulative probability functions or compare a numerically derived curves to a reference [31]. Here we compare the respective numerical univariate cumulative probability functions derived from synthetic samples with those derived from the sample. The difference metric, D_j_, for the KS test is the absolute maximum difference between the two cumulative probability functions under comparison.

*Mapping*: for each map in Step 1, we solve P_j_(x_j_) = G_j_(y_j_) numerically with interpolation methods described previously [32, 33], where P_j_(x_j_) and G_j_(y_j_) are assumed to be monotonically increasing. This solves the random variable transformation for each y_j_ analogous to histogram matching with double precision accuracy. Synthetic y_j_ (n = 10^6^) were generated as standardized normal random variables using the Box-Muller (BM) method. Maps from X to Y are expressed as y_j_ = m_j_(x_j_), where m_j_ is the j^th^ map. The corresponding inverse maps, 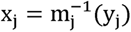, were derived numerically by inverting a given map and solving for x_j_. The map construction was complemented by generating synthetic x_j_ with Eq. [2] using h_j_ derived from DE optimization. Synthetic x_j_ generated here were not used further.

### 2.4 Synthetic population generation

Synthetic populations (SPs) were generated in the uncorrelated T representation and converted back to X via Y. In Step 2, the PCA transform for the sample is given by

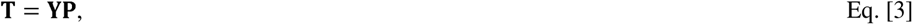

where **P** is a d×d matrix with uncorrelated normalized columns. These are the normalized eigenvectors of **C**_y_ that capture the sample’s covariance structure. **C**_t_ is diagonal with: 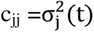 corresponding to the ordered eigenvalues of **C**_y_. We make the approximation that r(**t**) from the sample has the multivariate normal form expressed as

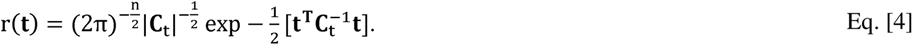

When the multivariate normality approximation holds in Y, it should hold in T. In Step 3, synthetic t_j_ were populated as zero mean independent normally distributed random variables with 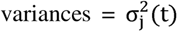 using the BM method, producing the SP in T (**T**_s_). The row length of T_s_ defines the number of realizations in each SP and is arbitrary; here we let n = 10^6^ for all SPs. In Step 4 to construct the SP in Y (**Y**_s_), the inverse PCA transform was used substituting **T**_s_ for **T** in Eq. [3] giving

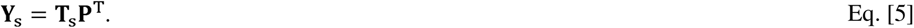

With this process, g(**y**) = g_n_(**y**) for synthetic data, and by premise the covariance of **Y**_s_ should be like that of **Y**. In Step 5 to produce the SP in X (**X**_s_), synthetic y_j_ were inverse mapped. Similarly, the covariance of **X** should be like that of **X**_s_. An example is also provided to illustrate that **X**_s_ is densely populated in contrast with the sparse sample in X.

### 2.5 Statistical methods

The goals are to evaluate the latent normal characteristic and to produce synthetic data that is statistically like its sample. This analysis is based on both multiple univariate/multivariate pdf and covariance comparisons. A given synthetic sample (n = 667) was drawn at random from its SP. The same realizations from a given synthetic sample were used for comparisons in X, Y, and T, when applicable.

#### 2.5.1 Probability density function comparisons

*Univariate pdf comparisons*: the KS test (described in Step 1) was used for all univariate pdf comparisons. For such comparisons, we selected the test threshold at the 5% significance level as the critical value. In X, p_j_(x_j_) from the sample were tested for normality and with their respective pdfs from synthetic samples from Step 5. In Y, g_j_(y_j_) from the sample were compared with the respective pdfs from their synthetic samples produced by Step 4; this implicitly evaluated univariate normality in Y because synthetic y_j_ were derived from a multivariate normal process in T_s_. In T, we compared r_j_(t_j_) from the sample with their respective pdfs derived from zero mean normally distributed random variables (i.e., synthetic t_j_) with 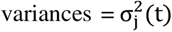 [from Step 3]. For each t_j_, the sample was compared with 1000 synthetic samples, and the percentage of times that measured D_j_ was less than the critical test value was tabulated.

*Distribution free multivariate pdf comparisons*: to evaluate whether the sample and synthetic samples were drawn from the same distribution without assumptions, we used the maximum mean discrepancy (MMD) [34] test. This is a kernel-based (normal kernel) analysis that computes the difference between every possible vector combination between and within two samples (excluding same vector comparisons). To determine the kernel parameter for these tests, we used the median heuristic [34, 35]. This analysis is based on the critical value (MMD_c_) at the 5% significance level and the test statistic 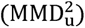. Both quantities are calculated from the two samples under comparison. This test has an acceptance region given the two distributions are the same: 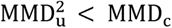 (see Theorem 10 and Corollary in [34]). This test was applied in X, Y, and T. Note, when applying this test in either T or Y, it is implicitly testing the sample’s likeness with multivariate normality. In X, Y, and T, 1000 synthetic samples were compared with the sample. The test acceptance percentage was tabulated. 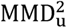 and MMD_C_ values are provided as averages over all trials because they change per comparison.

*Random projection multivariate normality evaluation*: random projections were used to develop a test for normality. The vector **w** with d components is multivariate normal if the scalar random variable, z = **u**^T^**w**, is univariate normal, where **u** is a d component vector with unit norm that is defined as a projection vector in this report [36, 37]. As motivated by Zhou and Saho [37], we developed this formulism into a specific random projection test. To actualize such a test to probe the samples and synthetic samples similarity with normality, the projection vector **u** was generated randomly 1000 times, referenced as **u**_s_. Here, s is the projection index ranging from [1,1000]. The projection equation is then expressed as

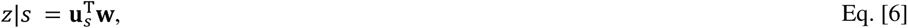

where z|s defines the scalar z conditioned upon s. In Eq. [6], **x**, **y**, or **t** was substituted for w in Eq. [6], and given projection was taken over all realizations (i.e., n = 667) of given sample. These realizations of z|s were used to form the normalized histogram that approximates the conditional pdf for the left side of Eq. [6] defined as f(z|s). A different series of **u**_s_ was produced for each representation; once a given series was produced, **u**_s_ remains fixed. The components of **u**_s_ were generated as standardized normal random variables, where **u**_s_ was normalized to unit norm. For a given sample, f(z|s) was tested for normality using the KS test. This procedure was repeated for all random projections (all s), resulting in 1000 KS test comparisons for normality. The percentage of the times that the null hypothesis was not rejected was tabulated as the normality similarity gauge; we refer to this as the *random projection test*, as it is the percentage that **w** was not rejected when probed in 1000 random directions. This test was performed once for each sample and with 100 synthetic samples and averaged. Here, the same **u**_s_ series used to probe a given sample was also used to probe 100 of its respective synthetic samples. We note, synthetic samples are multivariate normal in Y and T by their construction. Tests were performed on synthetic samples (Y and T) to give control standards as normal comparators. Tests were also performed in X as control comparator for the test itself and to determine if a given sample was multivariate normal before undergoing the mapping.

To gain insight into Eq. [6] and the test, expressions for both z|s and f(z|s) are developed. First, z|s results from a linear random variable transform given by

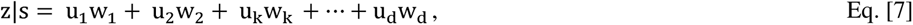

where u_k_ are the components of **u**_s_, and h_k_(w_k_) are the univariate pdfs for w_k_. To check one endpoint, we assume w_k_ are independent as a coarse approximation to our samples. Then, f(z|s) results from repeated convolutions given by

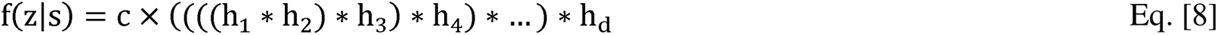

where c = u_1_×u_2_×u_3_× u_4_×…×u_d_, and h_k_(w_k_) ~ h_k_. If some h_k_(w_k_) have relatively much larger variances (widths) than others, their functional forms can tend to predominate Eq. [8].

*Mardia multivariate normality test*: this is a two component test that uses multivariate skewness and kurtosis for evaluating deviations from normality [36], applied in X, Y and T. It produces a deviation measure for each component as well as a combined measure; we cite the component-findings. This test was applied in X as a control. Outlier elimination techniques were not applied.

#### 2.5.2 Covariance comparisons

Two methods were used to evaluate the covariance similarity between samples and synthetic samples: with confidence intervals (CIs) and eigenvalue comparisons.

*Comparisons with confidence intervals*: each covariance matrix element between the sample and its respective synthetic samples were compared with CIs. We assumed the sample and synthetic samples were drawn from the same distributions. We used the elements from each **C**_x_ and **C**_y_ as point estimates from a given sample in both X and Y. One thousand synthetic samples (n = 667) were used to calculate 1000 covariance matrices (in X and Y). For each matrix element, the respective univariate pdf was formed, and 95% CIs were calculated. This procedure was repeated 1000 times. The percentage of times the sample’s point estimate (for each element in **C**_y_ and **C**_x_) was within the synthetic element’s CIs was tabulated.

*Comparisons with PCA*: the eigenvalues from **C**_y_ were used as the reference comparators under two conditions. For condition 1, the PCA transform determined with the sample was applied to a synthetic sample (sample/syn test); synthetic eigenvalues were estimated be calculating the variances of synthetic t_j_. For condition 2, the PCA transform determined with a synthetic sample selected a random was applied to the sample (syn/sample test); eigenvalues were estimated by calculating the variances of t_j_ from the sample. For both conditions, each eigenvalue (or equivalently, variance) was compared to its respective reference (sample) using the F-test.

## 3. Results

### 3.1 Univariate normality analysis in the X representation

Figures 1–3 show the univariate pdfs (solid) for each dataset in X for each sample, respectively. Of note, many pdfs are observably non-normal, usually right skewed. Each x_j_ in DS1 showed significant deviation from normality (p < 0.0001) except for x_9_. In DS2, neither x_1_ nor x_9_ (x_9_ is the same in DS1) showed significant deviations from normality, while the remaining x_j_ exhibited significant deviations (p < 0.0003) except for x_3_ (p = 0.0144). In DS3, x_1_ through x_5_ and x_9_ did not show significant deviations from normality, the remaining x_j_ deviated significantly (p < 0.002).

### 3.2 Mapping and KDE optimization

*Mapping*: Figure 5 shows an example of the X to Y map for y_9_ in the left-pane and the inverse Y to X map in the right-pane. Red-dashed lines show the map and its inverse constructed without synthetic x_j_, note the staircasing effects particularly in the tail regions, where sample densities are sparse. Black lines show the map and its inverse constructed with n = 667 (the sample) plus n = 10^6^ synthetic realizations produced with optimized uKDE. Note, the staircasing effects were removed when incorporating synthetic x_j_, which was common with all maps and inverses (not shown).

**Figure 5.**
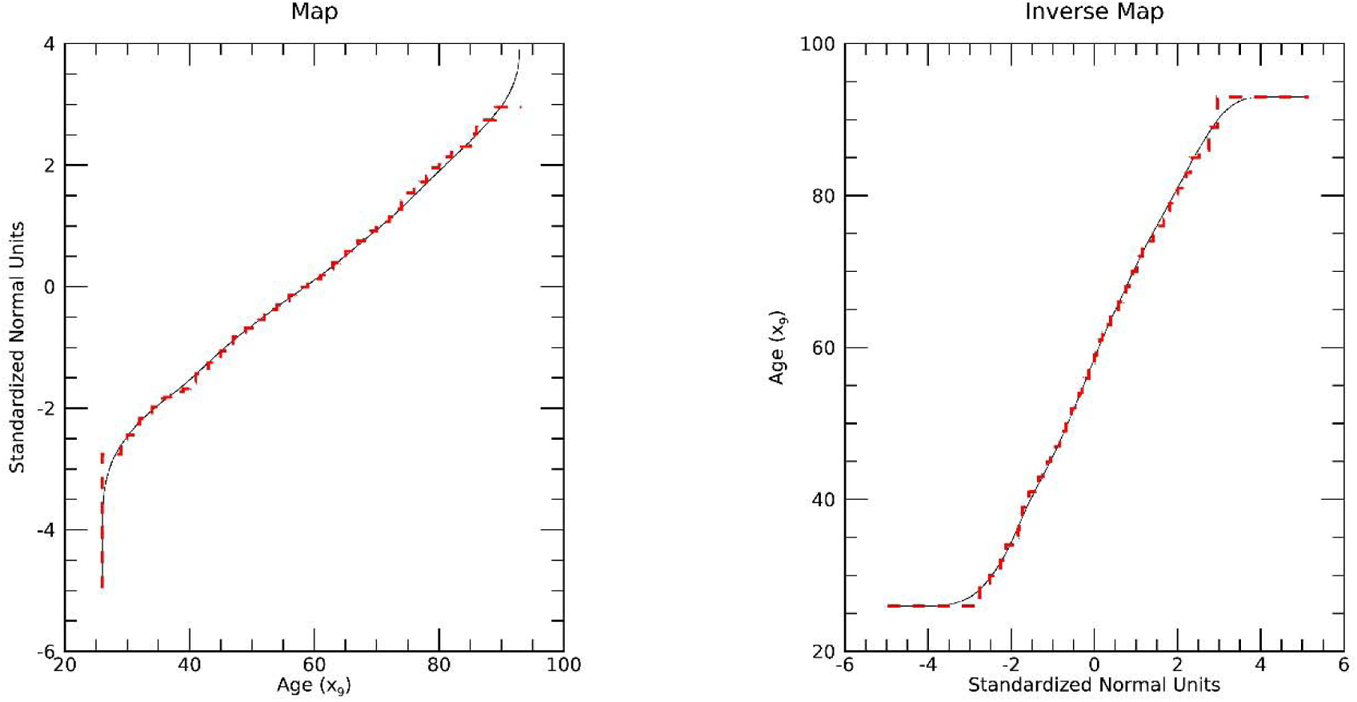
Univariate Mapping Illustration: this shows the map (left) and inverse (right) map for age (x_9_) in DS1 and DS2. Maps using the sample only (red-dashes) are compared with maps augmented with synthetic data (black-sold).

*uKDE optimization*: the optimization produced bandwidth parameter solutions (h_j_) used in Eq. [2]. Here, we illustrate the evolution of the *solution* with h_9_ from DS1 and DS2 as an example. Figure 6 shows the scatter plot between the candidate h_9_ population and the respective D_9_ (KS test difference metric) for DE generation = 1 in the left-pane and for the terminal generation = 30 in the middle-pane. Note, the solution space (middle-pane) is tightly clustered indicating DE *convergence*. A closer view of this cluster is shown in the right-pane of Figure 6. This relatively tight-cluster characteristic was common among all variables and datasets (not shown).

**Figure 6.**
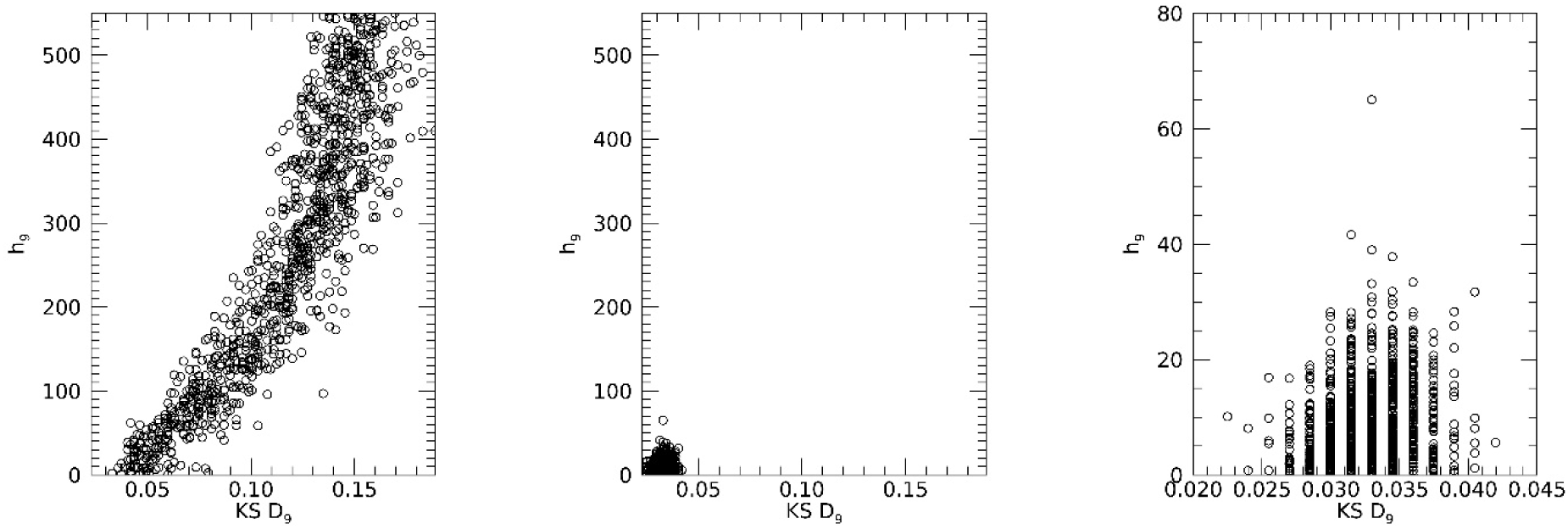
KDE Optimization Illustration: this shows the differential evolution optimization to determine h_9_ (for x_9_, age from DS1 and DS2). These show the scatter plots of the Kolmogorov Smirnov (KS) test metric (measured D_9_ quantities for entire generation) versus the h_9_ quantities for two generations: generation = 1 (left); terminal generation = 30 (middle); and closeup view of the terminal generation (right). The terminal generation shows the candidate solutions for h_9_ are tightly clustered (compare left pane with middle and right panes).

### 3.3 Univariate comparisons between samples and synthetic samples

*Comparisons in T*: these findings are summarized first because they start the flow back to X and can show departures from normality. Figures 7–9 show the pdfs for the samples (solid lines) compared with their corresponding synthetic pdfs (dashed). Table 5 shows the variances in T for each sample (i.e., eigenvalues for each sample). Due to (1) the normalization in Y, and (2) that d = 10, multiplying a given, 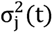 by 10% gives the percentage of the total variance explained by its t_j_. Table 6 shows the KS test findings for the univariate normality comparisons. Here we use a cutoff of < 65% to indicate deviation as most trends were well above this boundary. As shown in Table 6 (left column for each dataset), the normal model did not deviate for any t_j_ in DS1 (7 t_j_ were < 94%), deviated for t_10_ in DS2 (5 t_j_ were < 94%), and deviated for t_7_, t_8_, t_9_, and t_10_ in DS3 [3 t_j_ were < 94%]. In DS2, t_10_ explains about 0.2% of the total variance. Similarly, in DS3, the sum of the variances of the four variables (t_7_, t_8_, t_9_, and t_10_) constituted about 0.14% of the total variance.

**Figure 7.**
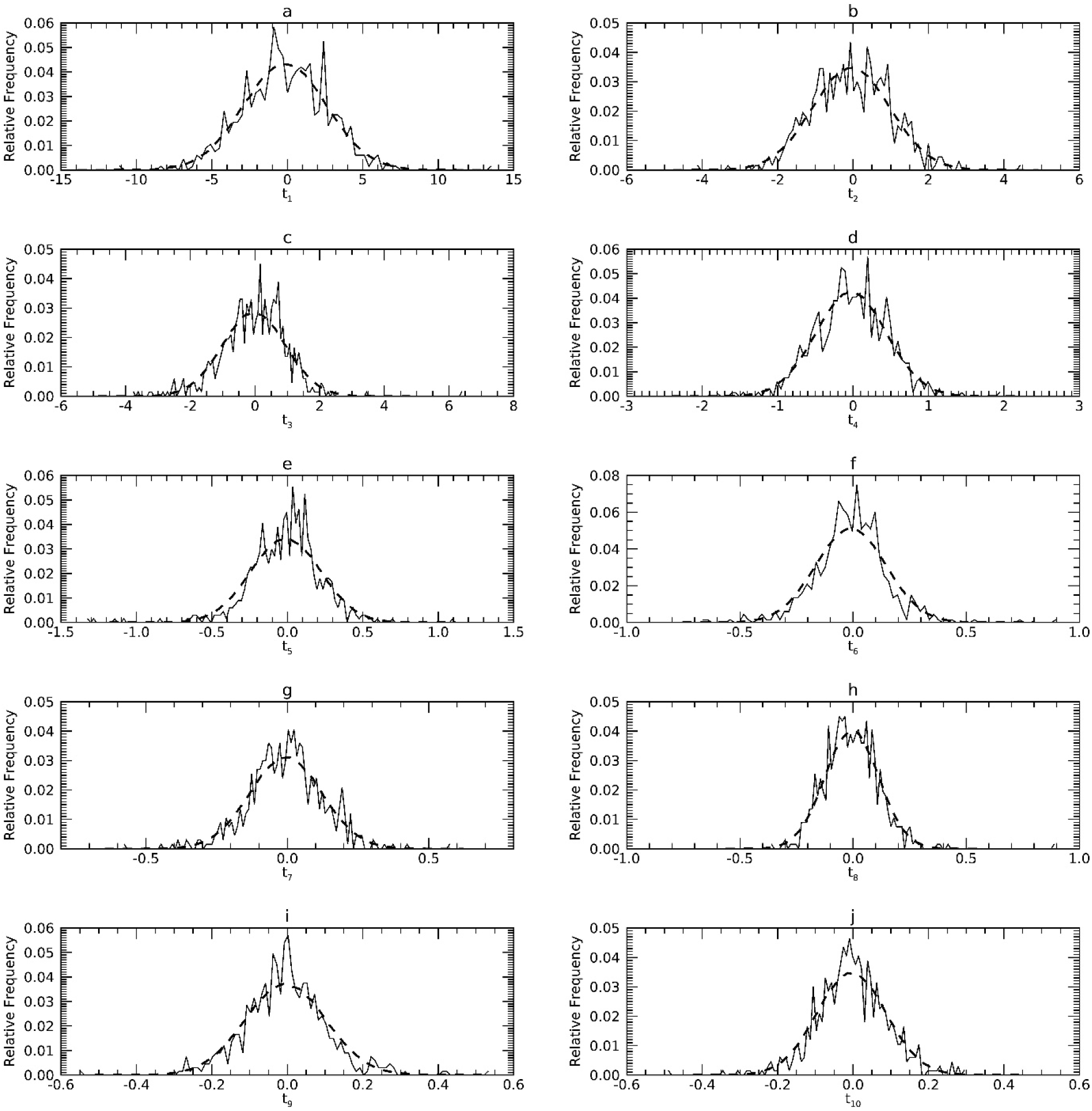
Marginal Probability Density Functions (pdfs) for DS1 in the T representation: each pdf for DS1 (sold) is compared with its corresponding pdf from synthetic data (dashes).

**Figure 8.**
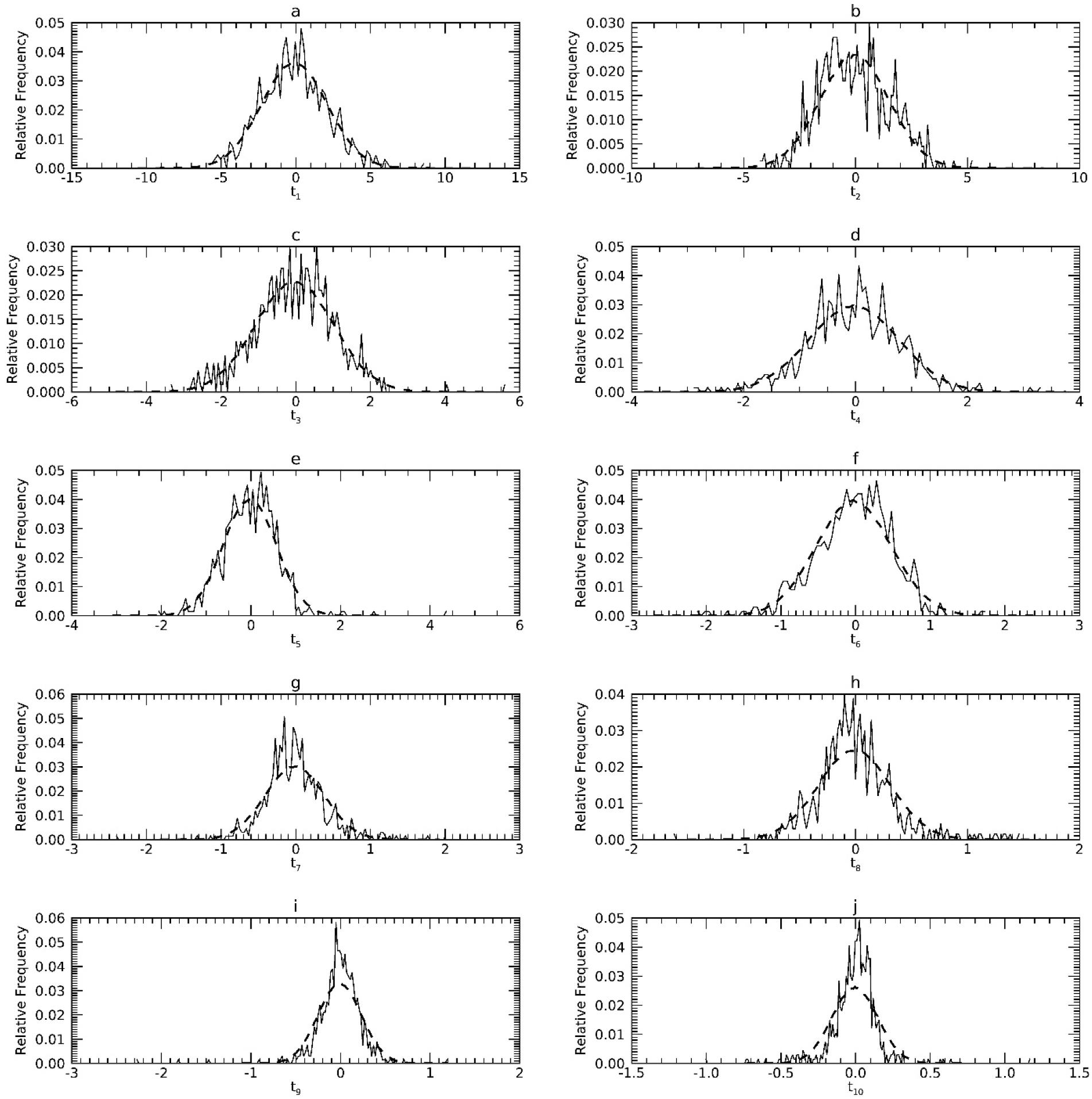
Marginal Probability Density Functions (pdfs) for DS2 in the T representation: each pdf for DS2 (solid) is compared with its corresponding pdf from synthetic data (dashes).

**Figure 9.**
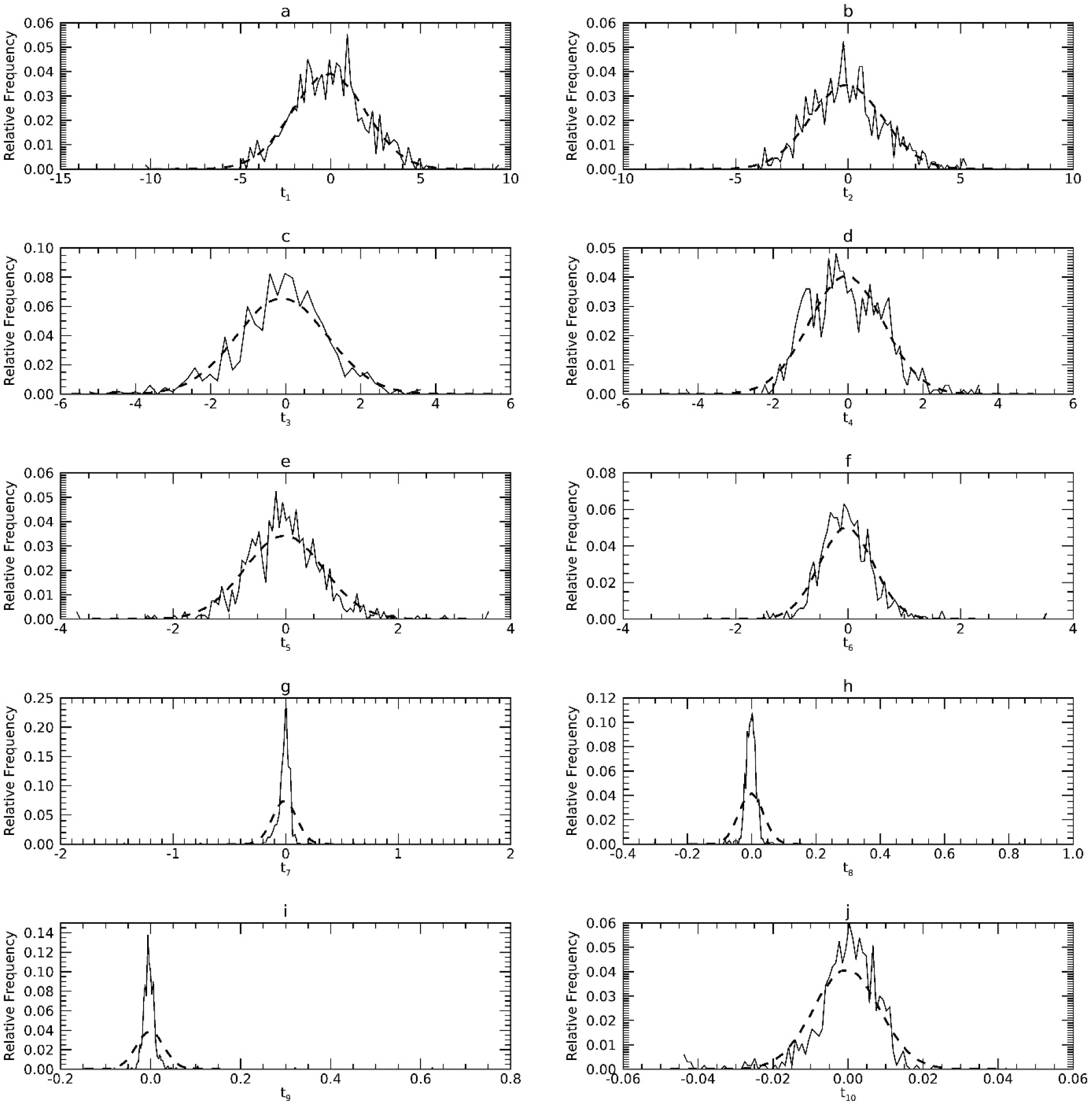
Marginal Probability Density Functions (pdfs) for DS3 in the T representation: each pdf for DS3 (solid) is compared with its corresponding pdf from synthetic data (dashes).

**Table 5.** Eigenvalue Comparisons. Within each dataset, the upper row (sample) gives the eigenvalues **C**_y_ (sample) used as the reference (ref) for comparisons. The Sample/Syn (condition 1) rows gives the eigenvalues (variances) determined by applying the PCA transform derived from the sample to a synthetic (Syn) sample. Syn/Sample (condition 2) rows give the eigenvalues (variances) determined by applying the PCA transform derived from **C**_y_ from a synthetic sample applied to the sample. An F-test was used to compare Sample/Syn t_j_ with the reference (ref) t_j_ and to compare Syn/Sample t_j_ with the reference t_j_. In all tests, the null hypothesis was not rejected (i.e. p > 0.05).

| | | $t_1$ | $t_2$ | $t_3$ | $t_4$ | $t_5$ | $t_6$ | $t_7$ | $t_8$ | $t_9$ | $t_{10}$ |
| --- | --- | --- | --- | --- | --- | --- | --- | --- | --- | --- | --- |
| DS1 | Sample (ref) | 7.6178 | 1.0682 | 0.9634 | 0.2208 | 0.0544 | 0.0244 | 0.0166 | 0.0142 | 0.0116 | 0.0085 |
|  | Sample/Syn | 7.6142 | 1.0700 | 0.9641 | 0.2213 | 0.0546 | 0.0244 | 0.0166 | 0.0145 | 0.0117 | 0.0087 |
|  | Syn/Sample | 7.6189 | 1.0987 | 0.9377 | 0.2183 | 0.0520 | 0.0246 | 0.0162 | 0.0138 | 0.0109 | 0.0090 |
| DS2 | Sample (ref) | 4.8873 | 2.3806 | 1.1083 | 0.6618 | 0.3575 | 0.2579 | 0.1576 | 0.1064 | 0.0590 | 0.0235 |
|  | Sample/Syn | 4.8857 | 2.3797 | 1.1072 | 0.6590 | 0.3565 | 0.2567 | 0.1647 | 0.1068 | 0.0596 | 0.0239 |
|  | Syn/Sample | 4.9237 | 2.2832 | 1.1094 | 0.6908 | 0.3691 | 0.2564 | 0.1714 | 0.1079 | 0.0634 | 0.0248 |
| DS3 | Sample (ref) | 4.1932 | 2.6215 | 1.4759 | 0.9790 | 0.4877 | 0.2286 | 0.0116 | 0.0015 | 0.0009 | 0.0001 |
|  | Sample/Syn | 4.1901 | 2.6125 | 1.4802 | 0.9837 | 0.4888 | 0.2294 | 0.0127 | 0.0015 | 0.0010 | 0.0001 |
|  | Syn/Sample | 4.2638 | 2.6817 | 1.3871 | 0.9368 | 0.4851 | 0.2312 | 0.0118 | 0.0014 | 0.0009 | 0.0001 |

**Table 6.**
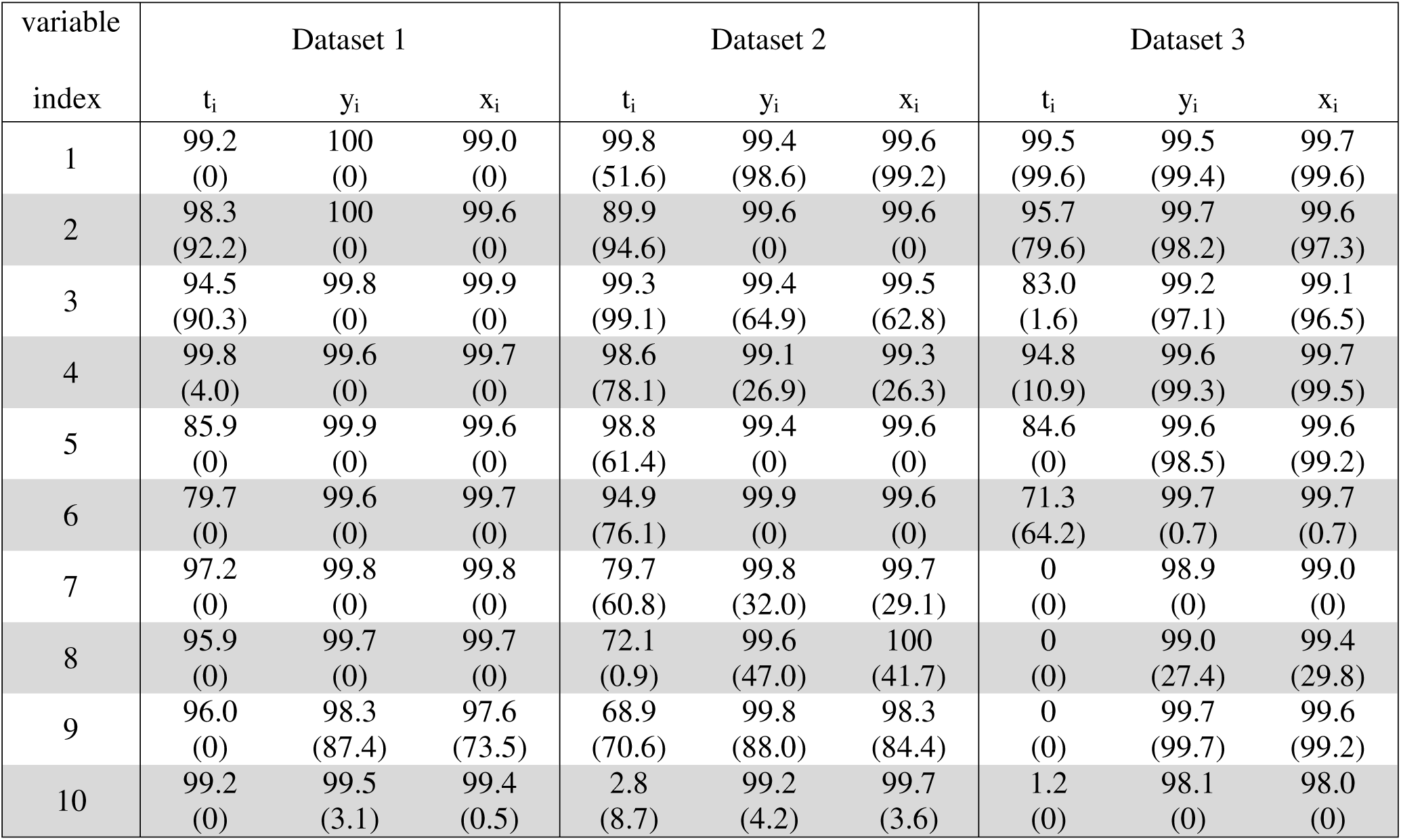
Univariate Kolmogorov Smirnov (KS) Tests: In each representation, probability density functions (pdfs) derived from synthetic samples were compared with the respective pdfs derived from sample for each variable. The KS test was applied 1000 times for each comparison. The number of times the null hypotheses was not rejected for a given test was tabulated as a percentage (all entries are percentages). Due to the experimental arrangements for y_j_ and t_j_, each test is equivalent to testing for normality as well. Parenthetical entries show the results when not using kernel density estimation to supplement the X-Y map constructions.

| variable | Dataset 1 |  |  | Dataset 2 |  |  | Dataset 3 |  |  |
| --- | --- | --- | --- | --- | --- | --- | --- | --- | --- |
| index | $t_i$ | $y_i$ | $x_i$ | $t_i$ | $y_i$ | $x_i$ | $t_i$ | $y_i$ | $x_i$ |
| 1 | 99.2<br>(0) | 100<br>(0) | 99.0<br>(0) | 99.8<br>(51.6) | 99.4<br>(98.6) | 99.6<br>(99.2) | 99.5<br>(99.6) | 99.5<br>(99.4) | 99.7<br>(99.6) |
| 2 | 98.3<br>(92.2) | 100<br>(0) | 99.6<br>(0) | 89.9<br>(94.6) | 99.6<br>(0) | 99.6<br>(0) | 95.7<br>(79.6) | 99.7<br>(98.2) | 99.6<br>(97.3) |
| 3 | 94.5<br>(90.3) | 99.8<br>(0) | 99.9<br>(0) | 99.3<br>(99.1) | 99.4<br>(64.9) | 99.5<br>(62.8) | 83.0<br>(1.6) | 99.2<br>(97.1) | 99.1<br>(96.5) |
| 4 | 99.8<br>(4.0) | 99.6<br>(0) | 99.7<br>(0) | 98.6<br>(78.1) | 99.1<br>(26.9) | 99.3<br>(26.3) | 94.8<br>(10.9) | 99.6<br>(99.3) | 99.7<br>(99.5) |
| 5 | 85.9<br>(0) | 99.9<br>(0) | 99.6<br>(0) | 98.8<br>(61.4) | 99.4<br>(0) | 99.6<br>(0) | 84.6<br>(0) | 99.6<br>(98.5) | 99.6<br>(99.2) |
| 6 | 79.7<br>(0) | 99.6<br>(0) | 99.7<br>(0) | 94.9<br>(76.1) | 99.9<br>(0) | 99.6<br>(0) | 71.3<br>(64.2) | 99.7<br>(0.7) | 99.7<br>(0.7) |
| 7 | 97.2<br>(0) | 99.8<br>(0) | 99.8<br>(0) | 79.7<br>(60.8) | 99.8<br>(32.0) | 99.7<br>(29.1) | 0<br>(0) | 98.9<br>(0) | 99.0<br>(0) |
| 8 | 95.9<br>(0) | 99.7<br>(0) | 99.7<br>(0) | 72.1<br>(0.9) | 99.6<br>(47.0) | 100<br>(41.7) | 0<br>(0) | 99.0<br>(27.4) | 99.4<br>(29.8) |
| 9 | 96.0<br>(0) | 98.3<br>(87.4) | 97.6<br>(73.5) | 68.9<br>(70.6) | 99.8<br>(88.0) | 98.3<br>(84.4) | 0<br>(0) | 99.7<br>(99.7) | 99.6<br>(99.2) |
| 10 | 99.2<br>(0) | 99.5<br>(3.1) | 99.4<br>(0.5) | 2.8<br>(8.7) | 99.2<br>(4.2) | 99.7<br>(3.6) | 1.2<br>(0) | 98.1<br>(0) | 98.0<br>(0) |

*Comparisons in Y*: figures 10–12 show the pdfs in Y resulting from the mapped samples (i.e., mapped x_i_) for each dataset (solid) compared with their respective synthetic pdfs (dashed), which are normal by construct. Comparisons in Y showed little departure from normality in any sample, as the tests were not rejected (< 99%) in most instances (Table 6, middle columns). Findings from DS2 and DS3 indicate that substituting normal pdfs in T whenever sample t_j_ deviated from normality had little influence on this analysis; this may be because these respective variables in total or isolation explained a minute portion of the variance in the respective PCA models.

**Figure 10.**
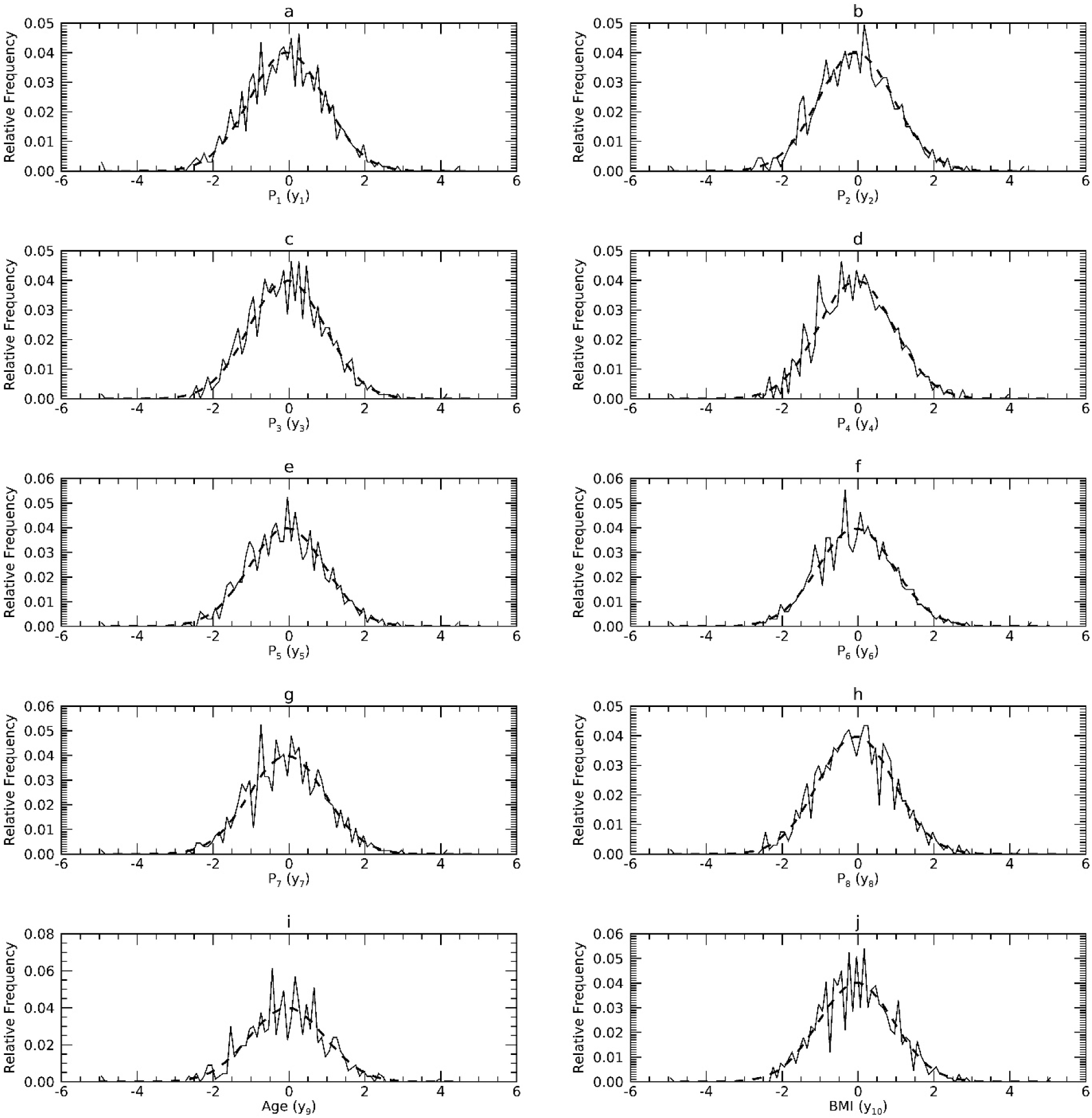
Marginal Probability Density Functions (pdfs) for DS1 in the Y representation: each pdf for DS1 (solid) is compared with its corresponding pdf from synthetic data (dashes).

**Figure 11.**
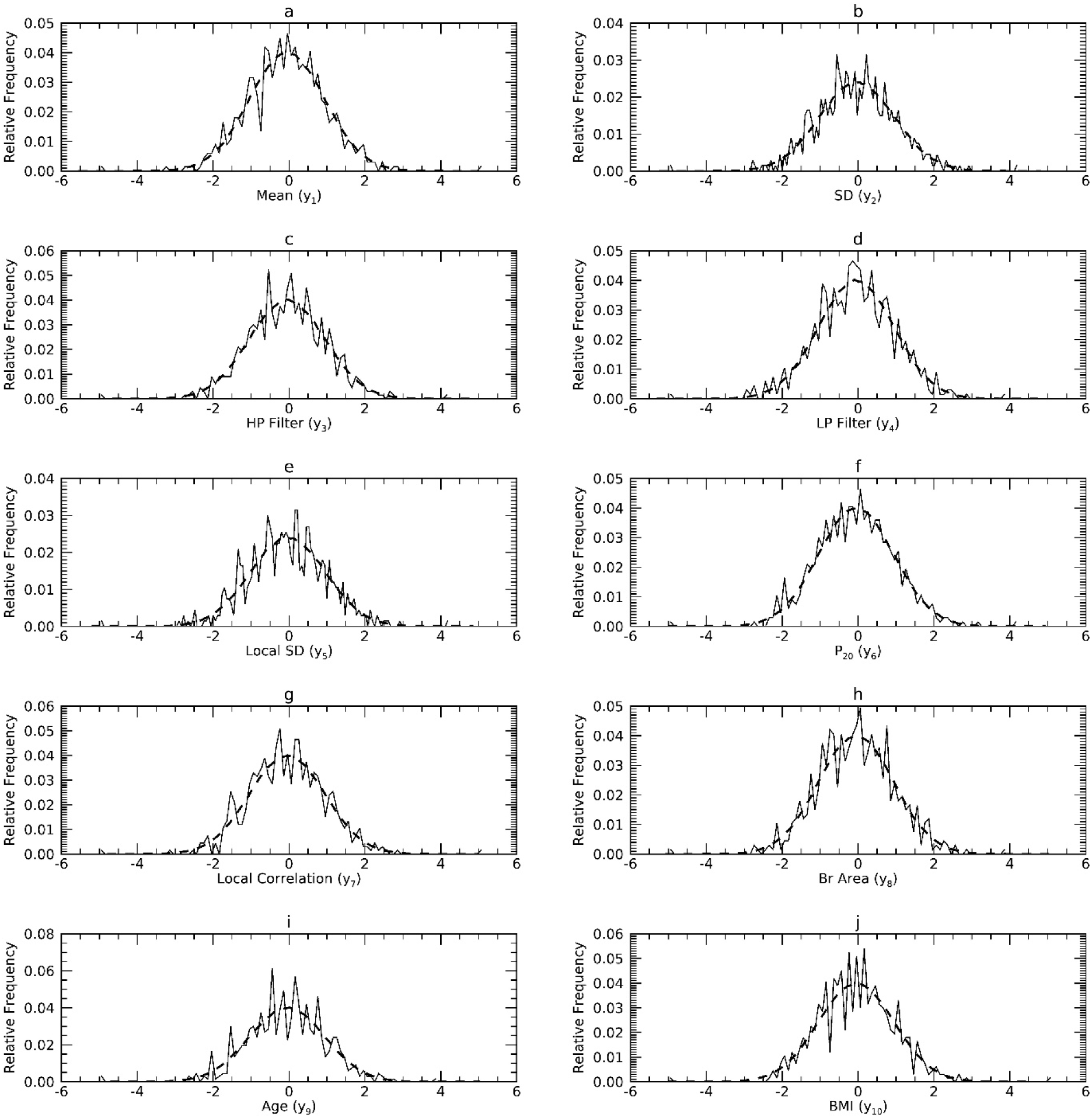
Marginal Probability Density Functions (pdfs) for DS2 in the Y representation: each pdf for DS2 (solid) is compared with its corresponding pdf from synthetic data (dashes).

**Figure 12.**
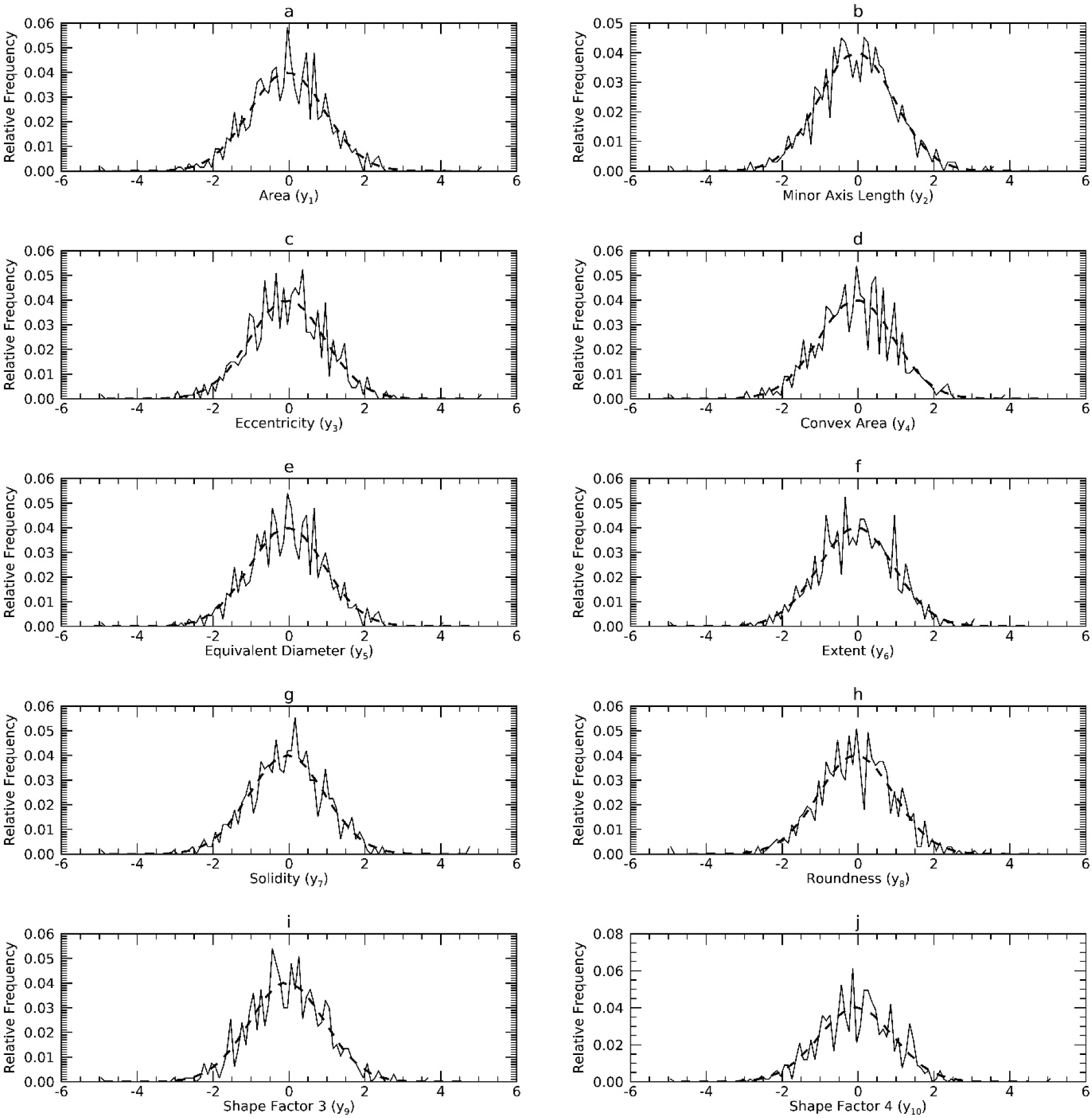
Marginal Probability Density Functions (pdfs) for DS3 in the Y representation: each pdf for DS3 (solid) is compared with its corresponding pdf from synthetic data (dashes).

*Comparisons in X*: figures 1–3 show the pdfs in X for the samples (solid) compared with their corresponding synthetic pdfs (dashed). The pdfs from the sample did not deviate from their corresponding synthetic pdfs in any dataset, as the tests were not rejected (< 99%) in most instances (Table 6, right columns). The parenthetical entries in Table 6 show the test findings without using synthetic data for the map/inverse constructions (using the samples only). These show that complementing the map constructions with uKDE is a necessary component of this methodology, although the degree of deviation from the KS tests varied across datasets. Note, the improvement held in D3 as well, which was normal in X (shown below).

### 3.4 Multivariate comparisons and normality comparisons

*MMD tests*: testing was performed in X, Y, and T, and the test metrics are provided in Table 7. These show the samples and respective synthetic samples were drawn from the same distributions. In 100% the tests, measured 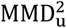 values were less than the critical MMD_c_ quantities. The MMD tests in Y and T were also proxy tests for sample-normality due to the SP constructs.

**Table 7.** Multivariate MMD Tests: The MMD test was used to compare the sample with 1000 synthetics samples. The MMD test quantities are averages over 1000 trials.

| MMD Test |  |  |  |
| --- | --- | --- | --- |
| **x** | DS 1 | $MMD_c$ | 0.2673 |
| | | $MMD_u^2$ | 0.0008 |
| | DS 2 | $MMD_c$ | 0.2667 |
| | | $MMD_u^2$ | -0.0002 |
| | DS 3 | $MMD_c$ | 0.2681 |
| | | $MMD_u^2$ | -0.0001 |
| **y** | DS 1 | $MMD_c$ | 0.2673 |
| | | $MMD_u^2$ | 0.0005 |
| | DS 2 | $MMD_c$ | 0.2667 |
| | | $MMD_u^2$ | -0.0008 |
| | DS 3 | $MMD_c$ | 0.2681 |
| | | $MMD_u^2$ | -0.0007 |
| **t** | DS 1 | $MMD_c$ | 0.2673 |
| | | $MMD_u^2$ | -0.0005 |
| | DS 2 | $MMD_c$ | 0.2667 |
| | | $MMD_u^2$ | -0.0006 |
| | DS 3 | $MMD_c$ | 0.2681 |
| | | $MMD_u^2$ | 0.0006 |

*Random Projection normality tests*: testing was performed in X, Y, and T, and the findings are shown in Table 8. Test findings were mixed for the samples in X and were not rejected for approximately these instances: 40% in DS1; 74% in DS2; and 99% in DS3. Thus, DS3 is better approximated as normal in X compared to the other samples. The tests for synthetic samples in X tracked the findings for their respective samples: 44%, 74% and 99% respectively. In Y, the tests for the samples were not rejected for about these instances: 99% in DS1, 97% in DS2, and 95% in DS3, whereas the test for the synthetic samples should no deviation from normality. Similarly in T, the tests for the samples were not rejected for about these instances: 99% in DS1, 94% in DS2, and 95% in DS3. There is a difference in the X and Y analyses because the mapping normalizes the variables in Y. In DS3, the standard deviations vary over many orders of magnitude (see Table 4). As shown by Eq. [8], variables in DS3 with the larger standard deviations may wash out the other variables; the variables in X that had normal marginals compared with those that were not, indicates that a portion of the normal marginals had much larger standard deviations (in Table 4, see x_1_-x_4_). As another control experiment, we standardized all variables in X to zero mean and unit variance and performed the tests again. The tests for the samples gave: 32.9%, 76.4 %, and 75.2% for DS1, DS2, and DS3, respectively. For synthetic samples, these tests gave: 33.6%, 80.1% and 89.1% for DS1, DS2, and DS3, respectively. Note, centering the means alone had no influence on the findings as expected (data not shown). Thus, normalizing the univariate measures can influence the likeliness with normality by virtue of Eq. [8]. In sum, these tests show all samples resemble multivariate normality in both Y and T and that the sample for DS3 resembles normality in X without mapping (without first normalizing the variances).

**Table 8.** Multivariate Normality Tests: For the random projection test, 1000 random projection vectors (1000 tests) were applied to a given sample and each test result was compared against normality; the percentage of times the test was not rejected was tabulated. For the synthetic (syn) data, the same 1000 projection vectors were used to test 100 synthetic samples from a given dataset. The percentage of times the test was not rejected was tabulated for each synthetic sample and percentages were averaged over the 100 synthetic samples. The Mardia tests was applied to the sample and to a synthetic sample selected at random.

|  |  |  | Random Projection | Mardia Skewness | Mardia Kurtosis |
| --- | --- | --- | --- | --- | --- |
| **x** | DS 1 | Sample | 40.2% | <0.0001 | <0.0001 |
|  |  | Syn | 41.2% | <0.0001 | <0.0001 |
|  | DS 2 | Sample | 74.3% | <0.0001 | <0.0001 |
|  |  | Syn | 74.1% | <0.0001 | <0.0001 |
|  | DS 3 | Sample | 98.9% | <0.0001 | <0.0001 |
|  |  | Syn | 99.8% | <0.0001 | <0.0001 |
| **y** | DS 1 | Sample | 98.9% | <0.0001 | <0.0001 |
|  |  | Syn | 100.0% | 0.9445 | 0.4287 |
|  | DS 2 | Sample | 97.4% | <0.0001 | <0.0001 |
|  |  | Syn | 99.9% | 0.4520 | 0.4374 |
|  | DS 3 | Sample | 94.8% | <0.0001 | <0.0001 |
|  |  | Syn | 100.0% | 0.2865 | 0.0859 |
| **t** | DS 1 | Sample | 99.0% | <0.0001 | <0.0001 |
|  |  | Syn | 100.0% | 0.1620 | 0.2143 |
|  | DS 2 | Sample | 94.1% | <0.0001 | <0.0001 |
|  |  | Syn | 100.0% | 0.8516 | 0.1933 |
|  | DS 3 | Sample | 94.5% | <0.0001 | <0.0001 |
|  |  | Syn | 100.0% | 0.6390 | 0.4010 |

*Mardia normality test*s: testing was performed in X, Y, and T. The findings are shown in Table 8. In X, the samples and synthetic samples all deviated from normality (both skewness and kurtosis). In either

Y or T, the samples showed significant deviations from normally in all tests. In contrast, synthetic samples did not deviate significantly from normality in any test in Y or T, as expected.

### 3.5 Covariance comparisons

*Covariance matrix comparisons with confidence intervals*: test findings are provided in tables 1–3 for the respective datasets. Part-a of each table shows the X quantities, and part-b shows the corresponding Y quantities. For DS1 (Table 1), covariance references (sample) were within the CIs of the synthetic data for 100% of the trials in both X and Y. In DS2 (Table 2), most references agreed with the synthetic elements except for two entries in X (Table 2a). From the 1000 trials, the x_2_x_3_ covariance was out of tolerance for 0.1% of the instances, and the x_5_x_10_ covariance was out for 18.7% of instances. For DS3, all covariance references were within tolerance except the x_7_x_10_ covariance, which was out of tolerance for 100% of the instances. In tests that showed more deviation (percentage > 0.1%), the reference covariances were approximately zero.

*Eigenvalue comparison tests*: eigenvalues are provided in Table 5, which is separated into three sections vertically. Reference eigenvalues are provided in the top row of each section. Eigenvalues calculated from the sample/syn (condition 1) and syn/sample (condition 2) are provided in the middle and bottom rows of each section, respectively. F-tests were not significant (p > 0.05) in any comparison with the references indicating similarity

### 3.6 Sample sparsity and synthetic population space filling

This illustration demonstrates that the approach fills in the multidimensional space with synthetic individuals derived from a relatively sparse sample. We selected a synthetic individual at random from DS1 giving this vector: **x**^T^ = [4.20, 2.08, 1.61, 1.15, 0.85, 0.67, 0.54, 0.44, 52.0, 23.6]. We selected x_1_ and x_8_ as the scatter plot variables: for the other 8 components, all synthetic individuals within x_ij_ ± ½ σ_j_ (the standard deviation for x_j_) were selected and viewed in the x_1_ - x_8_ plane as a scatter plot. The same vector and limits were used to select individuals from the sample. The plots are provided in Figure 13 for comparison. The sample (left-pane) produced n = 24 individuals, whereas the SP (right-pane) produced n = 36,398 individuals. These plots illustrate the sample’s relative sparsity and that the synthetic approach produces a dense population with individuals that did not exist in the sample.

**Figure 13.**
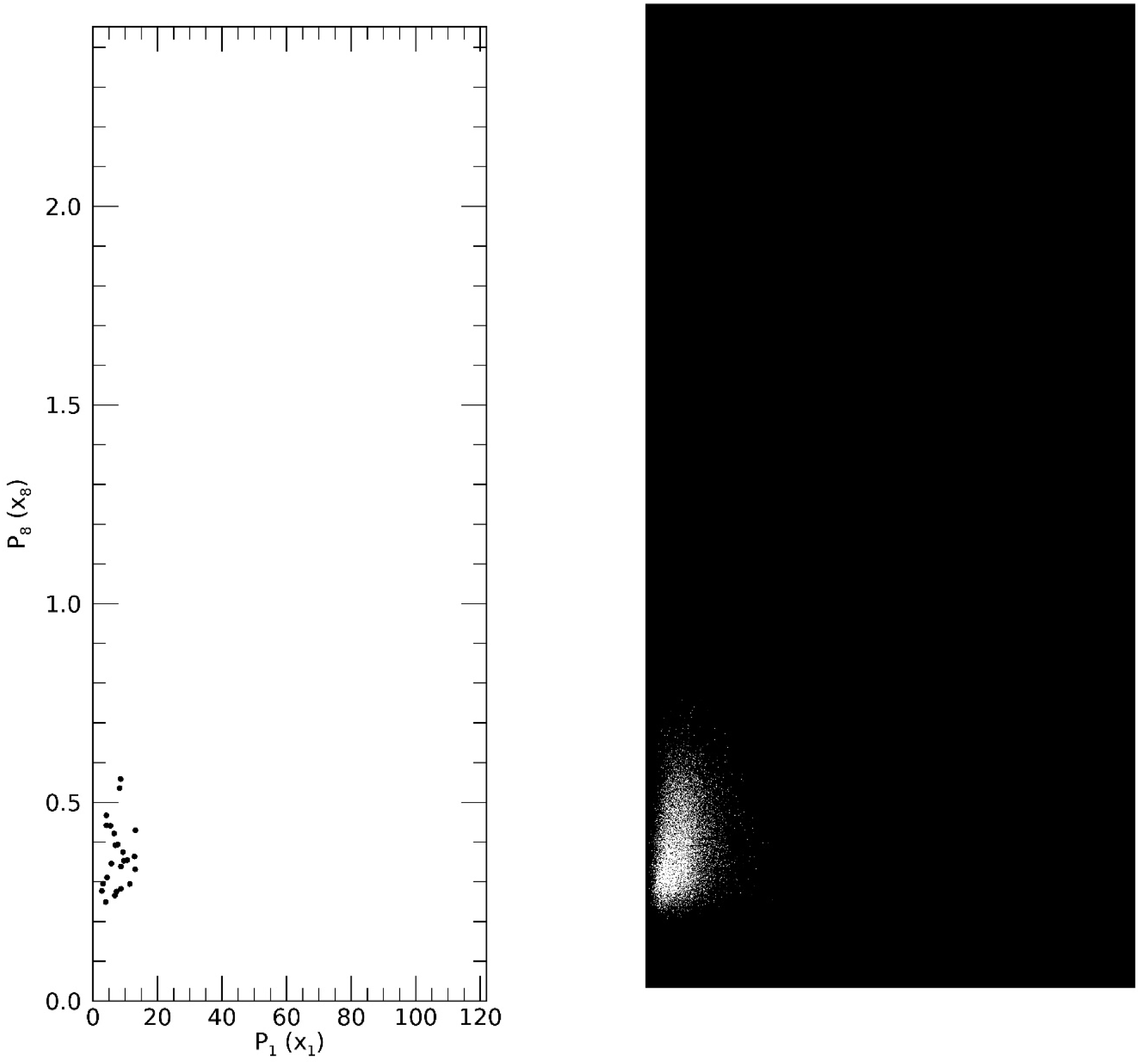
Sample and Synthetic Population Scatter Plot Illustration: A synthetic realization was selected randomly from DS1 giving this measurement vector: **x**^T^ = [4.20, 2.08, 1.61, 1.15, 0.85, 0.67, 0.54, 0.44, 52.0, 23.6]. x_1_ and x_8_ were selected for the scatter plot variables. For the other 7 components, all synthetic realizations within ± ½ (the respective standard deviation) were selected and viewed as a scatter plot in the x_1_x_8_ plane. Using the same vector and limits, individuals were selected from the sample in the same manner. The scatter plot for the sample (n = 24) is shown in the left-pane and from the synthetic population (n = 36,398) in the right-pane.

## 4. Discussion and Conclusions

The work involved several steps to generate synthetic data from arbitrarily distributed data. To the best of our knowledge, new aspects and findings from this work include: (1) demonstrating a class of arbitrarily distributed samples has a latent normal characteristic, as exhibited by two of the samples; (2) conditioning the input variables with sample size augmentation and univariate transforms so that known techniques could be applied to generate synthetic data; (3) deploying multiple statistical tests for assessing both normality and general similarity in the both the univariate and multivariate pdf settings; (4) developing methods for comparing covariance matrices; and (5) incorporating differential evolution (DE) optimization for uKDE bandwidth determination based on the KS fitness function. The related findings are discussed below in detail.

A method was presented that converts a given multivariate sample into multiple 1D marginal pdfs by constructing maps. These X-Y maps were constructed by augmenting the sample size with optimized uKDE. Performing the analysis with and without data augmentation improved the marginal pdf comparisons between the samples and synthetic samples; this also held in DS3 (four x_j_), which was approximately normal in X. PCA applied to standardized normal variables in Y produced uncorrelated variables in T, where synthetic data was generated. This approach essentially decouples the problem into the covariance relationships (in **P** and its inverse) and 1D marginal pdfs (i.e., approximate parametric models in Y and T). This decoupling is similar to the objective of Copula modeling that follows from Sklar’s work [38, 39]. Copula modeling allows specification of marginal pdfs and the correlation structure independently [40], noting that the marginals must be specified accurately and finding analytical solutions for d > 4 is difficult [41]. In contrast with Copula modeling, which is flexible, the covariance (or correlation) structure in our approach is fixed by the normal form and empirically derived; the marginals were forced to normality rather than specified. As a benefit, the CIs for **C**_k_ with our approach were estimated from the pdfs for each matrix element without assumption other than the normal calculation form. Additionally, the eigenvalue comparison technique results reinforced the CI comparison findings. However, it is not clear when (1) comparing the marginals with one set of tests, and (2) comparing the covariance relationships separately as another set of test results in a good overall empirical comparison-approximation between two multivariate samples, outside of the multivariate normal situation. Such situations will require further analyses.

There are several other points worth noting about this work. An empirically driven stochastic optimization technique was used to estimate the uKDE bandwidth parameters for the map/inverse constructions. The relative efficiency of the approach is an important attribute in that it only requires multiple uKDE applications rather than mKDE. The number of generations in the optimization was fixed. This can be changed easily to a variable termination based on achieving a critical threshold or applying other appropriate fitness functions. Likewise, there are plug-in kernel bandwidth parameters that can be used statically. These are derived by considering closed form expressions containing the constituent pdfs and minimizing the asymptotic behavior of either the mean integrated square error or mean squared error [42]. We explored such parameters [43], but they did not perform as well as the KS test with DE, notwithstanding the number of computations used here to determine a given bandwidth parameter. Of note, the KS test has limitations, as it is more sensitive to the median of the distribution rather than the tails. As an alternative, the Anderson Darling test is a variant of the KS procedure that is sensitive to the distribution tails [31]. The mapping from X to Y standardized the problem at the univariate level, but in general there is no guarantee that collectively it produced a multivariate normal in Y. Testing performed in Y (Step 2) could be used to discriminate input samples that have the latent normal characteristic from those that do not. The random project test could be developed into a gauge at this step for assessing the deviation from normality. Moreover, comparing r(t_j_) from the sample against normality (following Step 3) also provides a basis for testing sample’s likeness to a multivariate normal (discussed below). When p(**x**) is approximately multivariate normal, as in DS3, the mapping is not required and generating synthetic data based on PCA is a practiced technique; our approach addresses the case when this approximation fails to hold. The random projection tests changed the similarity with normality when standardizing the samples. Thus, the purposes for generating synthetic data should be considered before adjusting the input sample.

Several multivariate pdf tests were examined with mixed findings. MMD tests in X, Y, and T showed each sample was statistically similar with its respective synthetic sample(s). These tests also indicated normality in Y and T (by default). This MMD test is sensitive to changes in the mean. In our processing, all means were forced either to identically zero or to statistical similarity via mapping. Likewise, the heuristic used for the kernel bandwidth determination can be less than optimal under certain conditions, decreasing the test performance [44]. The random projection tests in Y and T indicated that the samples did not deviate from normality in most instances, whereas synthetic samples showed essentially no deviation. Understanding the acceptable departure from normality for this approach in the modeling context will require more work. This test also showed DS3 was approximately normal in X. In contrast, Mardia tests showed all samples deviated significantly from normality in X, Y and T, whereas synthetic samples showed essentially no deviation in Y or T as expected and significant deviation in X. Here, we made no attempt to mitigate possible outlier interference when analyzing the samples [45]. Note, testing for multivariate normality is not a trivial task; many of the complexities are covered by Farrell et al.[46].

The conclusions we make from these tests indicated each sample was approximately multivariate normal in Y and T, noting the approach may not be dependent upon this characteristic as elaborated below. In planned research, the worthiness of these approximations will be tested in the modeling context to evaluate whether sample and synthetic data are interchangeable. When this normality approximation holds, it implies that the original multivariate pdf estimation problem in X was converted to a parametric normal model described by Eq. [1], which simplifies the synthetic data generation. If this conversion generalizes to other datasets (at least in part), it implies that some class (subset) of the multivariate sample space can be studied with simulations by altering, n, d, and the covariance matrix to that of an arbitrary sample.

Alternatively, analyzing the samples in T may provide another method for comparing datasets, evaluating similarity, or generalizing our approach. The marginals from each sample were approximated as univariate normal in T, although there were noted variations. For example, and as noted, about 99% of the total variance came from the first four variables in DS1 (see Table 5). DS2 and DS3 were found to be similar with the first four variables accounting for 90-92% of the total variance. Thus, DS1 is more compressible than the other two datasets as expected due to the high correlation from its approximate functional Fourier form. Although not the purpose of this report, the amount of compression is a likely metric for estimating the effective dimensionality (d_e_) when d_e_ ≤ d, which could be useful for estimating sample size. When viewing the PCA transform through the NIPALS algorithm [14], it is clear when the total variance is explained by a number of components d_e_ << d, the remaining components are residue (noise, chatter, rounding errors). This effect could explain why DS2 and DS3 normality deviations in T did not influence the multivariate normal approximation in Y. Here, we did not encounter non-normal variables (from the samples) in T that explained a significant portion of the total variance. Future work will investigate causes for normality in T, i.e., possibly due to forced normality in Y, characteristics of the X representation data, or the PCA transform. If required, the technique could be generalized to accommodate non-normal marginals in T. As a generalization, uKDE will be investigated for generating univariate non-normal distributions in T when called for with the same method used to augment X for the map/inverse constructions. In this sample scenario, the Y description will deviate from the Eq. [1]. Although this premise will have to be investigated because the lack of correlation in T only guarantees t_j_ independence when r(**t**) is multivariate normal. We speculate when the sample has low correlation between most of the bivariate set in X, this approximation may hold.

Choosing the most appropriate space to perform modeling or to analyze the samples deserves consideration. We have used the covariance form suitable for normally distributed variables. In Y, this form is likely appropriate. We used the same form in X as well; this form may not be optimal here because covariance relationships are not preserved over non-linear transformations. It is our contention that Y is best suited for modeling because the marginals are normal. It is common practice in univariate/multivariate modeling to adjust variables (univariately) to a standardized normal form or apply transforms to remove skewness. The X to Y map converted each x_j_ to unit variance; when the natural variation for x_j_ is important, the mapping can be modified easily to preserve the variance. If the variable interpretation is not important, modeling can also be performed in T.

The method in this report addresses the small sample problem given the sample has the latent normal characteristic or is normal. The approach will require further evaluation on different datasets to understand its general applicability and when the univariate mapping from X to Y approximately produces a multivariate normal. Multiple methods were explored to evaluate multivariate normality. These tests indicated that the samples approximated normality in both the Y and T but also showed some deviation from normality. The interpretation of these findings in the context of data modeling may aid in understanding the limits of both the multivariate SPs and normality approximations in this report’s data and beyond. For example, determining the limiting percentage of the random projection tests may be informative in the modeling context. In summary, we offer a definition for an insufficient sample size in the context of synthetic data. When considering a given sample with d attributes and specified covariance structure, a sample size that does not allow reconstructing its population can be considered as insufficient. In future work, we will apply the methods in this report to understand the minimum sample size, relative to d and a given covariance structure, that permits recovering the population.

## 5. Declarations and Compliance with Ethical Standards

*Funding*: The work was in part supported by Moffitt Cancer Center grant #17032001 (Miles for Moffitt) and National Institutes of Health Grants R01CA166269 and U01CA200464.

*Competing interests*: The authors have no competing interests to declare that are relevant to the content of this article.

*Ethics and consent to participate*: Mammography data was collected under protocol #Ame13_104715, Institutional Review Board (IRB), University of South Florida, Tampa, FL.

*Consent of publication*: The work does not contain personal identifiers.

*Availability of data*: The link to publicly available data is provided in the text. Mammography summary data can be obtained upon request. Kernel parameters are also available upon request.

*Author contributions*: JH is the corresponding author, conceived the plan and methods; EF is a coauthor, developed the computer code and assisted in the plan and methods development; AB is a coauthor and provided statistical and principal component analysis expertise; MJS is a coauthor and provided statistical expertise; SE is a coauthor and assisted in the plan and methods developments.

## Notes

### Competing Interest Statement

The authors have declared no competing interest.

### Summary of Updates

Version 2: Techniques to Produce and Evaluate Realistic Multivariate Synthetic Data

